# Integrated transcriptome, DNA methylome and chromatin state accessibility landscapes reveal regulators of Atlantic salmon maturation

**DOI:** 10.1101/2020.08.28.272286

**Authors:** Amin R. Mohamed, Marina Naval-Sanchez, Moira Menzies, Bradley Evans, Harry King, Antonio Reverter, James W. Kijas

## Abstract

Despite sexual development being ubiquitous to vertebrates, the epigenetic mechanisms controlling this fundamental transition remain largely undocumented in many organisms. Through whole-methylome, whole-transcriptome and chromatin landscape sequencing, we discovered global control mechanisms as well as specific regulators of sexual maturation in Atlantic salmon. This large integrated study was based on an experimental time course that successfully sampled the period when Atlantic salmon commence their trajectory towards sexual maturation. Co-analysis of DNA methylome and gene expression changes revealed chromatin remodelling genes *arid1b* and *smarca2* were both significantly hypermethylated and upregulated in the ovary during the onset of maturation. We also observed changes in chromatin state landscape occurred early in the transition and were strongly correlated with fundamental remodelling of gene expression. Finally, we integrated our multiomics datasets to identify *trim25* and *znf423* as key regulators in the pituitary that underwent 60 fold change in connectivity during the transition to sexual maturation. The study provides a comprehensive view of the spatiotemporal changes involved in a complex trait and opens the door to future efforts aiming to manipulate puberty in an economically important aquaculture species.

## Main

Epigenetic regulation of gene expression influences a vast spectrum of complex traits, with examples spanning the onset and severity of human disease, developmental transitions during growth and the expression of ecologically and economically relevant traits across the animal kingdom. Our understanding of the epigenetic contributions to trait variation remains low in comparison to causative genes derived from approaches such as genome wide association studies. However, the relatively recent development of sophisticated sequence-based assays for the detection of chromatin state changes and methylation status have enabled the landmark development of genome wide maps of regulatory elements in human ^1,2,3^, mouse ^4,5,6^ and other model organisms such as the fruit fly (*Drosophila melanogaster*) and the nematode (*Caenorhabditis elegans*) ^7,8^. These have provided the impetus for a plethora of research focussed on understanding epigenetic mechanisms and their role regulating gene expression.

Sexual maturation is a fundamental transition ubiquitous to vertebrates and provides a model for the study of epigenetic regulation. The key tissues are known given the reproductive cycle is regulated by activation across the brain, pituitary, gonadal (BPG) axis in organisms spanning mammals to teleost fish. Further, upon maturation these tissues undergo known and often profound transcriptomic remodelling providing a large dynamic range to increase the likelihood of identifying regulatory networks ^9,10,11,12^. The genetic architecture of sexual maturation has been extensively studied using association studies in both wild and farmed Atlantic salmon populations ^13,14,15,16^. Despite the importance of sexual maturation as a trait of interest it can be difficult to study, as the timing of onset varies widely in response to both genetics and environmental factors and occurs prior to measurable phenotypic change. To overcome this, we chose to investigate sexual maturation in Atlantic salmon where photoperiod manipulation in an experimental system can be used to synchronise animals and access tissues across the time period when animals first commit to the onset of puberty. We also chose a multiomics approach, which has the power to identify the control mechanisms underpinning complex traits ^17,18^. We describe changes in gene expression, DNA methylation and chromatin accessibility to identify epigenetic mechanisms associated with maturation in a commercially important aquaculture species.

## Results

### Initiation of Atlantic salmon sexual maturation and a multiomic workflow

Fish were managed in a tank based experimental system to facilitate a long-light photoperiod regime known to stimulate the onset of sexual maturation (Fig. 1a) ^19,20^. Fish from a single management group were sacrificed at a timepoint immediately before initiation of the long-light regime (throughout referred to as T1) and at three timepoints afterwards (T2, T3 and T4). An increase in gonadal somatic index (GSI) of sampled fish across the time course confirmed an active response to the long photoperiod (Fig. 1b). Significant increases were observed only at T4 (t-test P-value = 0.021). Tissues from the brain – pituitary – gonad axis and liver were sampled at each timepoint to form the basis of a multiomics workflow for data generation and integrative analysis spanning the transcriptome, DNA methylome and chromatin state datatypes (Fig. 1c; Supplementary Fig. S1).

**Fig.1.**
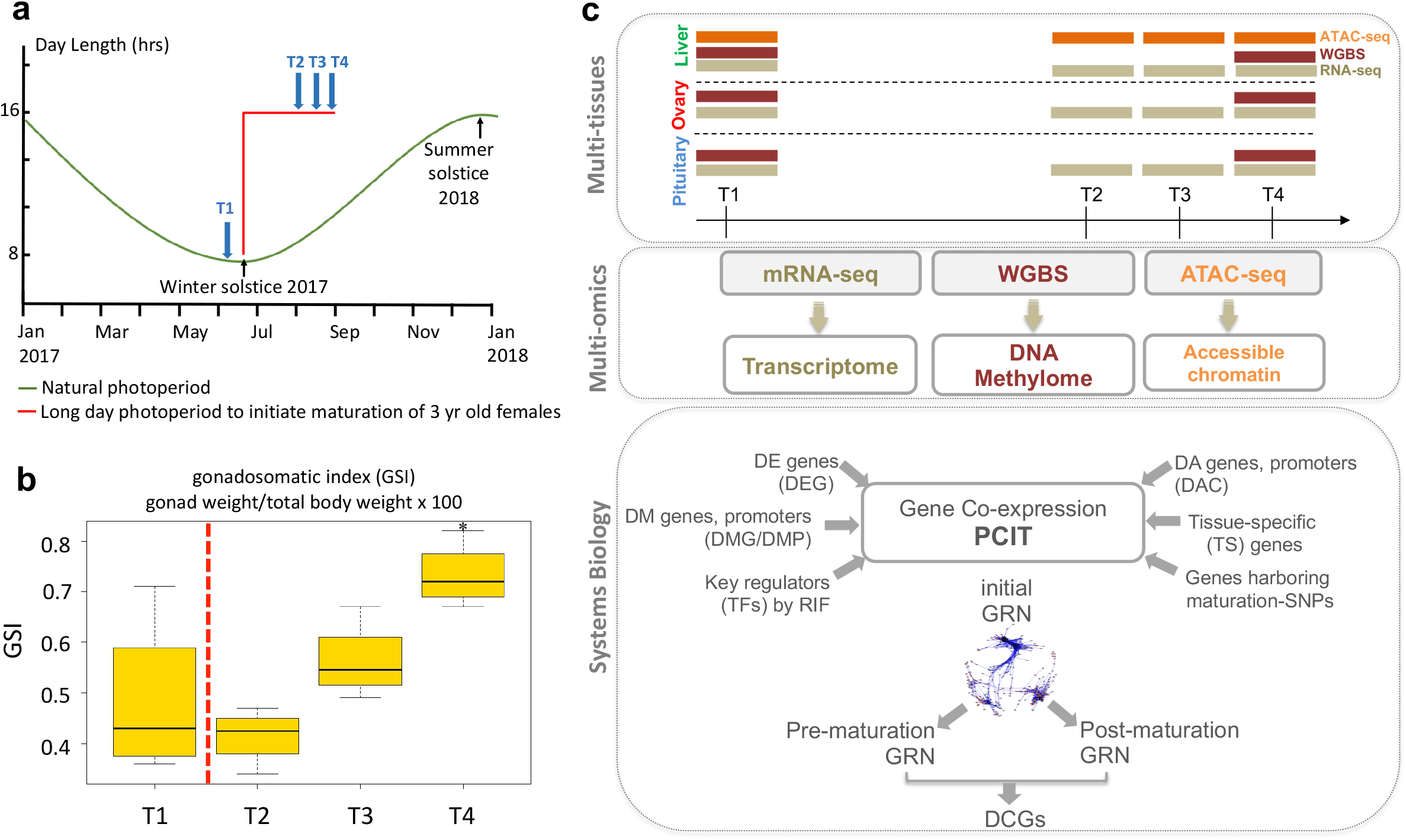
Multi-tissue transcriptomic and epigenomic changes associated with onset of salmon maturation. **a, Induction of maturation through photoperiod manipulation and sampling time points.** Animals were managed via photoperiod manipulation to synchronise the timing of commitment into maturation. 4 fish were sampled at each of the T1 (before long day photoperiod signal) and T2-T4 time points (during maturation) in 2 weeks intervals to control for variation between individuals. **b, Gonadosomatic index (GSI) throughout the time-course experiment**. GSI increased gradually from T2 till the last sampling event at T4 indicating active response towards maturation in these animals. The variability in GSI measures during maturation decreased compared to that of T1. **c, Tissue collection, multi-omics integrated analyses and gene regulatory networks (GRNs)**. Samples from the pituitary gland, ovary and liver were collected at each sampling event. High-throughput sequencing was utilised to profile genome-wide changes in transcriptomes, DNA methylomes in the three tissues along with chromatin accessibility in liver. These genome-wide results were integrated with knowledge of key regulators (TFs) and other results from previous work that identified maturation associated-SNPs and tissue-specific genes. PCIT algorithm was then utilised to construct gene regulatory networks (GRNs). Pre- and post-maturation GRNs were constructed to identify differentially connected genes (DCGs).

### Maturation leads to significant transcriptome changes

We sequenced messenger RNA (mRNA) from four biological replicates of each tissue before and after the onset of maturation. A total of 3.2 billion paired-end reads were mapped against the Atlantic salmon reference genome with 72% mapping efficiency to create an average depth of 50 million reads per library (Supplementary Table 1). Consistency across biological replicates within each timepoint was high for each tissue except for brain (see Supplementary Results; Supplementary Fig. S2). To begin characterization of transcriptomic remodelling, we compared gene expression levels in three post-maturation samples (T2, T3 and T4) to the control T1 (Fig. 2a). The pituitary gland showed comparatively few transcriptomic responses that involved 543 differential expressed genes (DEGs) (Supplementary Table S2). In contrast, more widespread remodelling was observed in both ovary (5,993 DEGs) and liver (9,541 DEGs, adjusted *P* < 0.05 (Fig. 2a, Supplementary Tables S3, S4). The number of DEGs increased with elapsed time following the onset of the long light photoperiod for the two BPG axis tissues (pituitary and ovary). Of these, the ovary underwent the most dramatic remodelling over time with 403, 1,709 and then 3,497 DEGs observed at timepoints T2, T3 and T4 respectively. This increasing trajectory of differential gene expression, coupled with the elevated GSI following the light stimuli (Fig. 1b), strongly suggests the experimental approach successfully initiated the onset of maturation.

**Fig.2.**
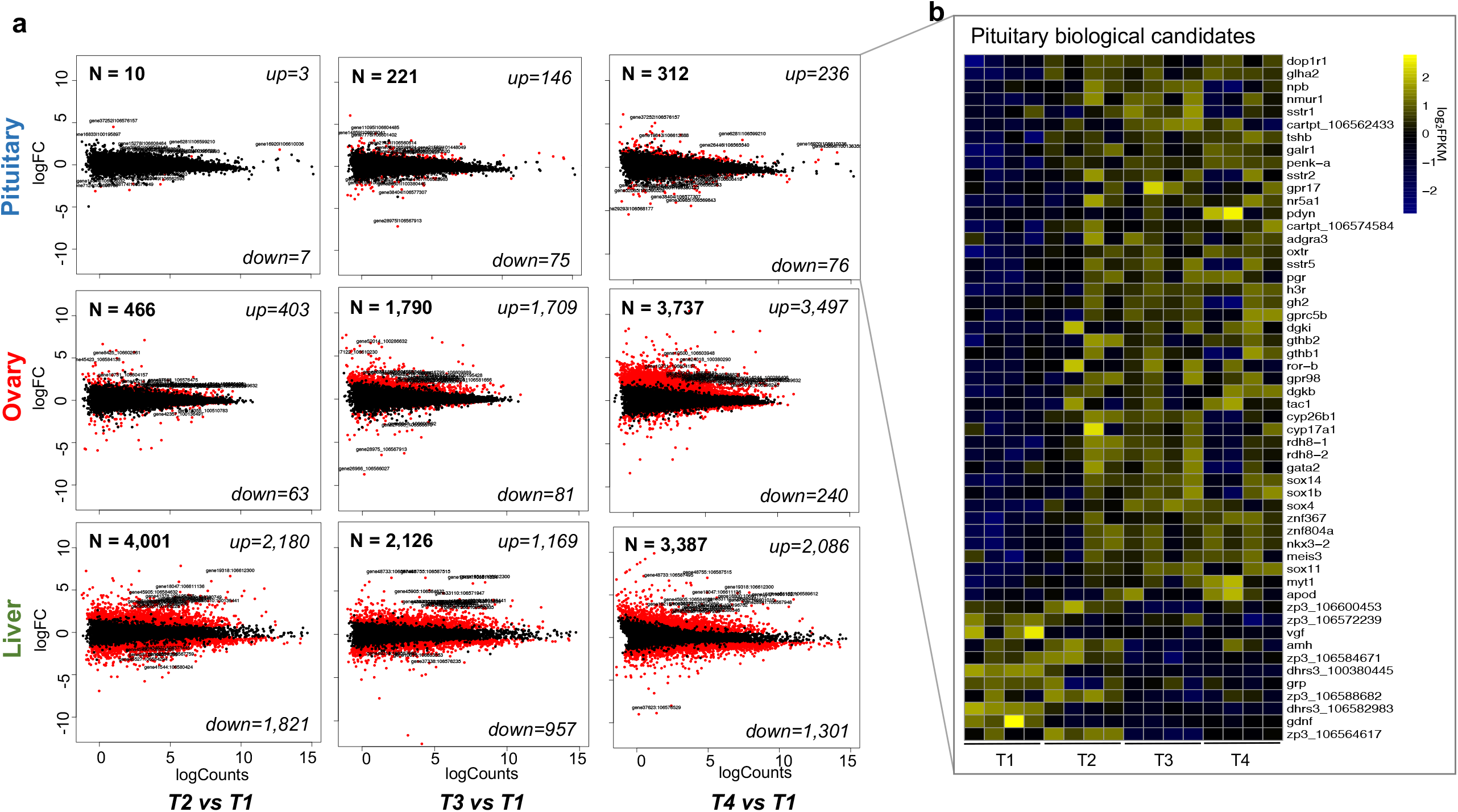
Significant transcriptome remodelling driving onset of salmon maturation. **a**, 9 MA plots showing differentially expressed genes (DEGs) in pituitary, ovary and liver (FDR < 0.05 and log2fold change >± 1) at T2, T3 and T4 during maturation compared to control samples at T1. Transcriptomic results reveal gene upregulation in both pituitary gland and ovary during maturation. **b**, Heatmap of pituitary candidate genes with significant expression. The hierarchical clustering was obtained by comparing the log2-transformed and mean-centred FPKM values. Upregulated genes in the pituitary include several genes with reproduction-related functions such as steroidogenesis (*cyp17a1*), hormone receptors (*oxtr, pgr, dop1r1, galr1*), genes coding for pituitary hormones (*gh2, gthb1, gthb2, glha2*), retinoic acid (RA) signalling (*cyp26b1, rhd8-1/2)*, sex-related TFs (several sox genes and *gata2*). The upregulation of gonadotropins subunits (*gthb1, gthb2* and *glha2*) is highly significant in the context of maturation onset. *gthb1* and *gthb2* encoding gonadotropin subunits beta-1 and 2 and *glha2* encoding glycoprotein hormones alpha chain that constitutes the salmon gonadotropins (*gth-1* and 2).

### Upregulation of pituitary hormones

The pituitary is expected to play a key role in the early triggers for maturation onset (21). Hierarchical clustering of the pituitary DEGs revealed two distinct groups, one of upregulated genes (n=333; 60% of pituitary DEGs) and one of downregulated genes (n=125). The upregulated group showed significant Gene Ontology (GO) enrichment for maturation-related functions including *G protein−coupled receptor signalling* and *hormone activity* (Supplementary Fig. S4 and Supplementary Table S5). Pituitary DEGs were involved in several reproduction-related functions such as steroidogenesis (*cyp17a1*), hormone receptors (*oxtr, pgr, dop1r1, galr1*), genes coding for pituitary hormones (*gh2, gthb1, gthb2, glha2*), retinoic acid (RA) signalling (*cyp26b1, rhd8-1*/*2*) and sex-related transcription factors (TFs) (several sox genes and *gata2*). The upregulation of gonadotropins subunits is highly significant (Fig. 2b). This included both *gthb1* and *gthb2* that encode the gonadotropin subunits beta-1 and 2 as well as *glha2* that encodes the glycoprotein hormone alpha chain. Together, these form the heterodimeric gonadotropins *gth-1* and *gth-*2 that have previously been shown to stimulate gonadal growth in the juvenile stages of both rainbow trout and coho/chum salmon ^22,23,24^. Further, physicochemical characterization of the salmon gonadotropins indicate they are functionally related to follicle stimulating hormone (fsh) and luteinizing hormone (lh) in vertebrates (reviewed by ^25^). We find *glha2* (the common subunit present in gonadotropins) was consistently upregulated in the pituitary throughout the experiment (Supplementary Fig. S3). Our results directly confirm the action of these key pituitary hormones in the onset of maturation. To characterize the transcriptomic remodelling occurring in the ovary and liver, we assessed the DEG sets for GO enrichment. Upregulated genes in the ovary revealed processes related to *cell adhesion, immune/inflammatory response* and *development* (Supplementary Fig. S5; Supplementary Table S6), while gene families involved in *organic acid metabolic processes* and *mitochondrial transport* were enriched among liver upregulated genes (Supplementary Fig. S7 and Supplementary Table 7). Maturation-related functions including steroidogenesis, hormonal receptors and follicular development were identified in ovary (see Supplementary Results; Supplementary Fig. S6).

### DNA methylome maps of three Atlantic salmon tissues

To investigate the regulatory mechanisms controlling differential expression, the first genome-wide CpG methylation maps were developed for Atlantic salmon using whole-genome bisulfite sequencing (WGBS). Methylome data was collected from two biological replicates at the terminal time points (T1 and T4, Fig. 1c) from three tissues (pituitary, ovary and liver), generating 2.6 billion paired-end uniquely mapped reads (average coverage of 11x) (Supplementary Table S8). We found a genome-wide methylation rate of 81% per sample (Supplementary Fig. S8; Supplementary Table S8), which is similar to the rate observed in vertebrate genomes (60-90%) ^26,27^. Methylome data was assessed based on coverage, read mapping and consistency between biological replicates (see Supplementary Results; Supplementary Table S8; Supplementary Fig. S8c, d). Among the different dinucleotide contexts, CpG methylation contributed the vast majority (∼ 99.5% on average) compared to CHH or CHG methylation which were excluded from further analysis (Supplementary Fig. S8a; Supplementary Table S8).

By comparing the CpG methylation patterns between T4 and T1 samples (Fig. 3a), we identified 1902, 2982 and 1606 differentially methylated regions (DMRs) in the pituitary gland, ovary and liver respectively (Fig. 3b; Supplementary Fig. 9a, b; Supplementary Tables 9, 10, 11). The average length of DMRs was short (251 bp) and their distribution was both genome wide (Fig. 3b) and highly non-random, with 52% found to overlap protein coding genes and another 18% located within 5 kb upstream or downstream (Fig. 3c; Supplementary Fig. 9c). The location of DMRs was also strongly tissue specific, with few regions shared between tissue pairs and only 8 found in all three tissues (Supplementary Fig. 9b). Next, we investigated the directionality of DMRs across tissues and found approximately equal rates of hyper-methylation (increased methylation in T4) and hypo-methylation (decreased methylation in T4) in both the pituitary and liver. Strikingly, the majority of DMRs in ovary were hyper-methylated (2175 DMRs or 73%), independent of their genomic location (genic regions, promotors, 5 kb downstream or intergenic) (Supplementary Fig. 9d). This is consistent with DNA methylation having important roles during epigenomic reprograming in embryo and stem cells development ^28^, compared to highly stable methylomes in somatic cells ^29^. This pronounced skew towards increased methylation occurred in the tissue with both the highest total number of observed DMRs and the largest increase in upregulation of gene expression. Of the tissues investigated, the ovary also undergoes the most radical physiological change during maturation as it transforms via vitellogenesis and oocyte development in preparation for egg release during spawning.

**Fig.3.**
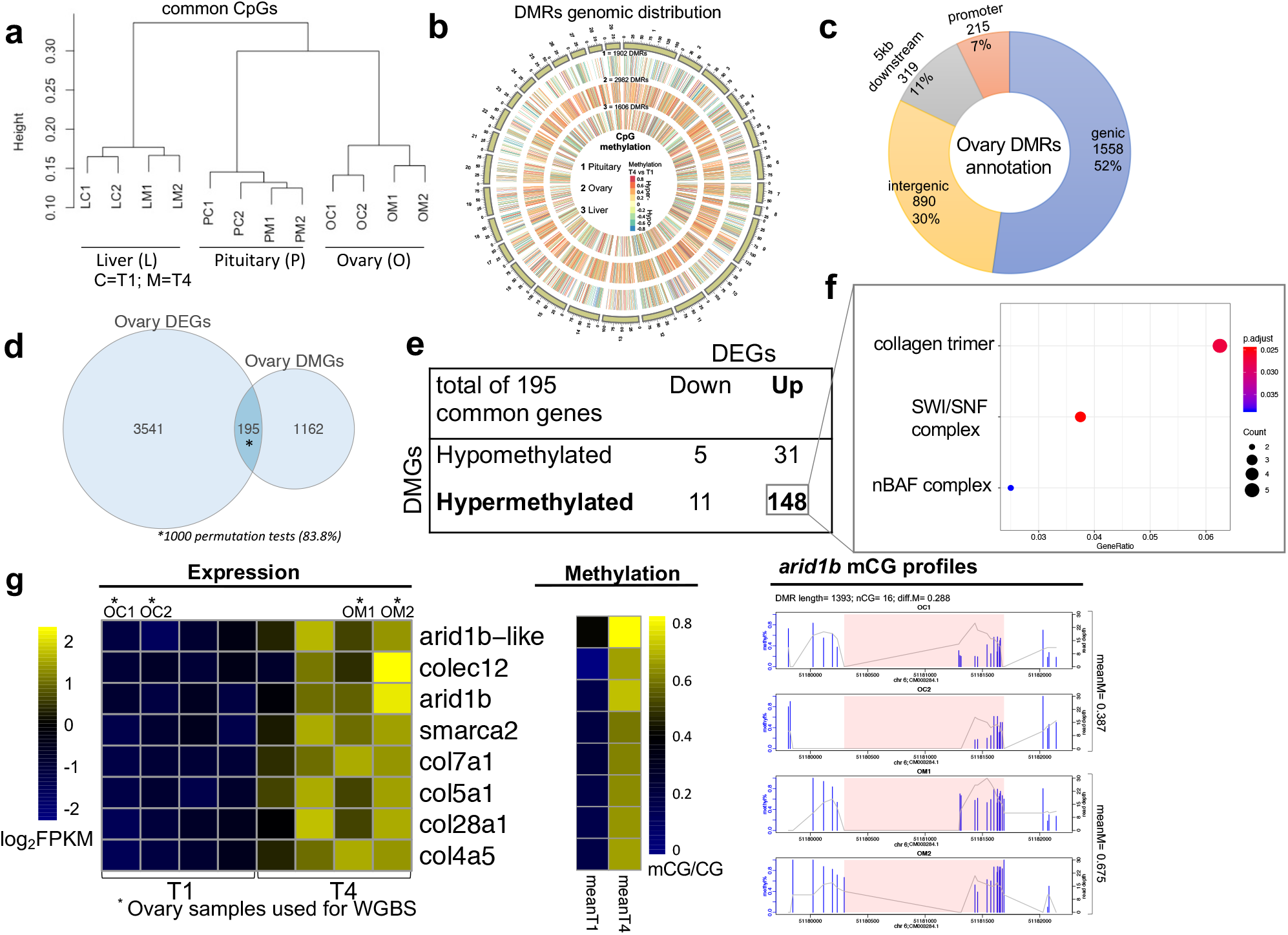
Multi-tissue DNA methylome maps reveal methylated genes with key roles in salmon maturation. **a**, Hierarchical clustering of the common CpGs in the 12 methylome libraries reveals clustering by tissue type and grouping pituitary and ovary in a cluster and liver in the other. This shows variation in methylation is much lower between replicates of the same sample compared with variation among tissues as expected for high quality data. **b**, Circos plot shows genome-wide distribution of the significant differentially methylated regions (DMRs) detected in T4 samples compared to the control T1 in pituitary, ovary and liver (from the outer circle inwards). **c**, Genomic distribution of ovary DMRs across gene models in the salmon genome shows the majority co-located within genes, promoter defined as genomic regions located 5kb upstream of transcription start sites (TSSs). **d**, Significant overlap between ovary DMGs and DEGs. **e**, DNA methylation and expression directionality at gene bodies confirms that gene body methylation positively affects gene expression. **f**, Enriched gene ontology (GO) terms (hypergeometric test, Bonferroni-adjusted *P* < 0.05) among the list of 148 hypermethylated genes in ovary. **g**, Heatmaps showing significant upregulation of genes driving GO enrichment shown in part f, mean CpG methylation levels of upregulated genes before and after maturations and CpG methylation profiles for the *arid1b* gene. The methylation plot shows percentages of methylated CpGs along with coverage depth at each CpG site. Pink rectangles represent the differentially methylated regions. Genomic coordinates are indicated below the density plot and DMR details are also indicated.

### Differentially methylated genes serve key roles in maturation

The gene catalogue present within differentially methylated regions (DMRs) was assessed for their function in relation to the trait. The majority of differentially methylated genes (DMGs) in ovary were hypermethylated (n=1165; 74%) (Supplementary Fig. S9d) and significantly enriched for three biological process (GO-BP), one cellular component (GO-CC) and 24 molecular function (GO-MF) terms including those with maturation-related functions such as *semaphorin/glutamate receptor activity* (Supplementary Fig. S10; Supplementary Table S12). Among these genes, at least 33 genes have demonstrated roles in the biology of maturation including follicular development (*plexnb1, sema4f* coding for Plexin-B1 and Semaphorin-4f) ^30^ and the control of gonadotropin-releasing hormone excitability (*grm8* coding for the glutamate receptor 8) ^31^. We plotted the normalised expression of these candidates from samples collected at T1 and T4 and found that the majority were upregulated at T4, consistent with their hyper-methylated status (Supplementary Fig. S11).

### Co-analysis of DNA methylome and transcriptome reveals role for chromatin remodelling during maturation

The identification of significant transcriptional and methylation changes allowed us to explore the dynamic between these two processes by assessing the overlap of genes declared as both DEG and DMG. The overlap was low and non-significant for liver (38 / 616 or 6% of DMGs were also DEGs) and pituitary (11 / 762 or 1.4% of DMGs were DEGs) (Supplementary Fig. S9e). However, 195 or 14% of ovary DMGs (195 / 1357) were also differentially expressed, a number that exceeded random expectation in 83.8% of 1000 permutations tests (Fig. 3d). This suggests changes in methylation status may directly control gene expression in this subset of genes. If true, we would expect to see correspondence between the directionality of the expression and methylation changes. This appeared to be the case, as 82% of upregulated genes (148 / 179; Binomial P-value = 8.727E-20) were hyper-methylated at T4 relative to T1 (Fig. 3e), matching the classical expectation of gene body methylation mediated control of gene expression ^32,33^. The 148 genes were enriched for 3 GO-CC terms related to chromatin remodelling complexes (SWI/SNF and nBAF) (Fig. 3f; Supplementary Table S13), and the associated genes displaying coordinated expression and methylation status (Fig. 3g). For example, the *arid1b* gene encodes AT-rich interactive domain-containing protein 1B and *smarca2* encodes the global transcription activator snf2l2. Both proteins are involved in chromatin remodelling as they are core components of the SWI/SNF remodelling complexes^34^ that carry out enzymatic change to chromatin structure by altering DNA-histone contacts ^35^. This opens the possibility they act as key control points, to regulate a wide array of other genes during ovarian development. A final examination was performed to search for evidence of a generalised and genome wide association between gene expression levels and methylation status. Expression for all genes with either differential gene body or promoter methylation is shown in Supplementary Fig. S12. We found no correlation for either comparison, consistent with previous studies that reported weak correlation between DNA methylation and gene expression in humans ^36,37^ and more recently in fish ^38^. Taken together, the results confirmed that while methylation alone does not control genome-wide patterns of gene expression, it plays a key role upregulating a defined set of genes during the maturation process.

### Early and stable changes in the chromatin state

To deepen the characterization of the epigenomic features during maturation, we performed ATAC-seq (assay for transposase-accessible chromatin sequencing^39^) to produce genome-wide maps of chromatin accessibility changes. ATAC-seq was performed for multiple tissues and peak enrichment around transcription start sites used as the key quality control metric (TSS, Supplementary Figure S13). Following data pruning, we took 12 liver libraries (3 replicates across all 4 timepoints) and a total of 699 million uniquely mapped paired-end reads (Supplementary Table S14) forward into joint analysis with RNA-seq and WGBS data. Principal component analysis (PCA) of the 12 ATAC-seq samples revealed the T1 replicates formed a tight cluster positioned separately from the T2 – T4 samples, which were less well distinguished from each other (Fig. 4a). The first two principal components accounted for 50% of variation, which is less than the comparative analysis of the same tissue using RNA-seq (66% Fig. 4a). To characterise changes in chromatin state following long light initiation, we defined differentially accessible regions (DARs) where mapping counts differed significantly between T1 and other time points. This revealed a strong early remodelling in the chromatin state landscape, as most DARs were observed at T2 (n=1501) before decreasing in stepwise fashion at T3 (n=477) and T4 (n=148; Fig.4b; Supplementary Table S15). The direction of change was approximately balanced between DARs with increased and decreased accessibility, broadly matching the balance between up and down regulated global gene expression changes observed for liver (Fig. 2a). We next asked if the early changes in chromatin state persisted throughout the time course using hierarchical clustering. The majority of DARs (n=1036 or 57%) exhibit reduced accessibility at T2 compared with T1 and subsequently remained unchanged at later time points (Fig. 5a; Supplementary Figure S14). Similarly, regions that gained accessibility at T2 (n=696 or 38%) also remained unchanged at later timepoints. This left less than 10% of DARs (n=99) that displayed an oscillating pattern following the onset of the maturation. Together, this revealed the ATAC-seq signatures were predominantly stable chromatin state changes, as opposed to pulsatile epigenomic changes that snapped back after a small number of days or weeks.

**Fig.4.**
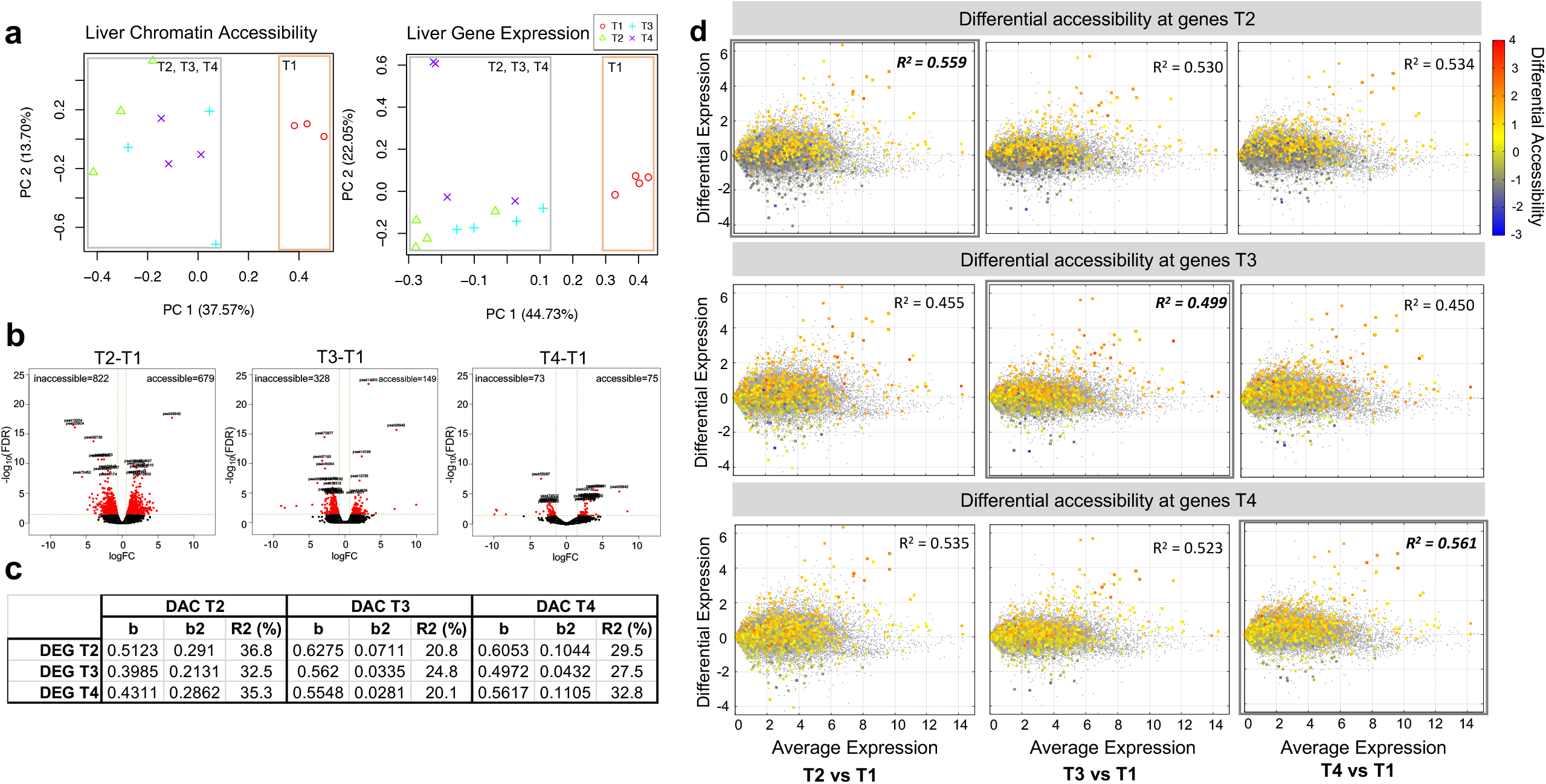
Genome-wide maps of chromatin accessibility and gene expression changes. **a, Combined ATAC-seq and RNA-seq analysis.** Principal component analysis (PCA) conducted using normalised (log2CPM) values of the lists of significant differentially accessible regions (DARs) and significant differentially expressed genes (DEGs) at T2, T3 and T4 during maturation compared to control samples at T1, the same significance thresholds were applied (FDR < 0.05 and log2fold change >±1). **b, Strong and early remodelling in the chromatin state landscape**, the volcano plots show differentially accessible regions (DARs) where mapping counts differed significantly between T1 and other time points. **c, Regression analysis conducted on significant DARs located at gene bodies and the corresponding gene expression data**. The table shows that accessibility at T2 explains the majority of the observed differential expression throughout the experiment. **d, Chromatin accessibility and gene expression are positively correlated**. 9 MA biplots showing genome-wide gene expression and overlain differentially accessible regions dynamics at gene bodies in liver at T2, T3 and T4 compared to T1. Chromatin accessibility levels are shown in red-blue spectrum reflecting open-to closed chromatin at gene bodies and the corresponding gene expression in grey colour. Note that R^2^ values are the highest when using gene expression and accessibility from the same time point.

**Fig.5.**
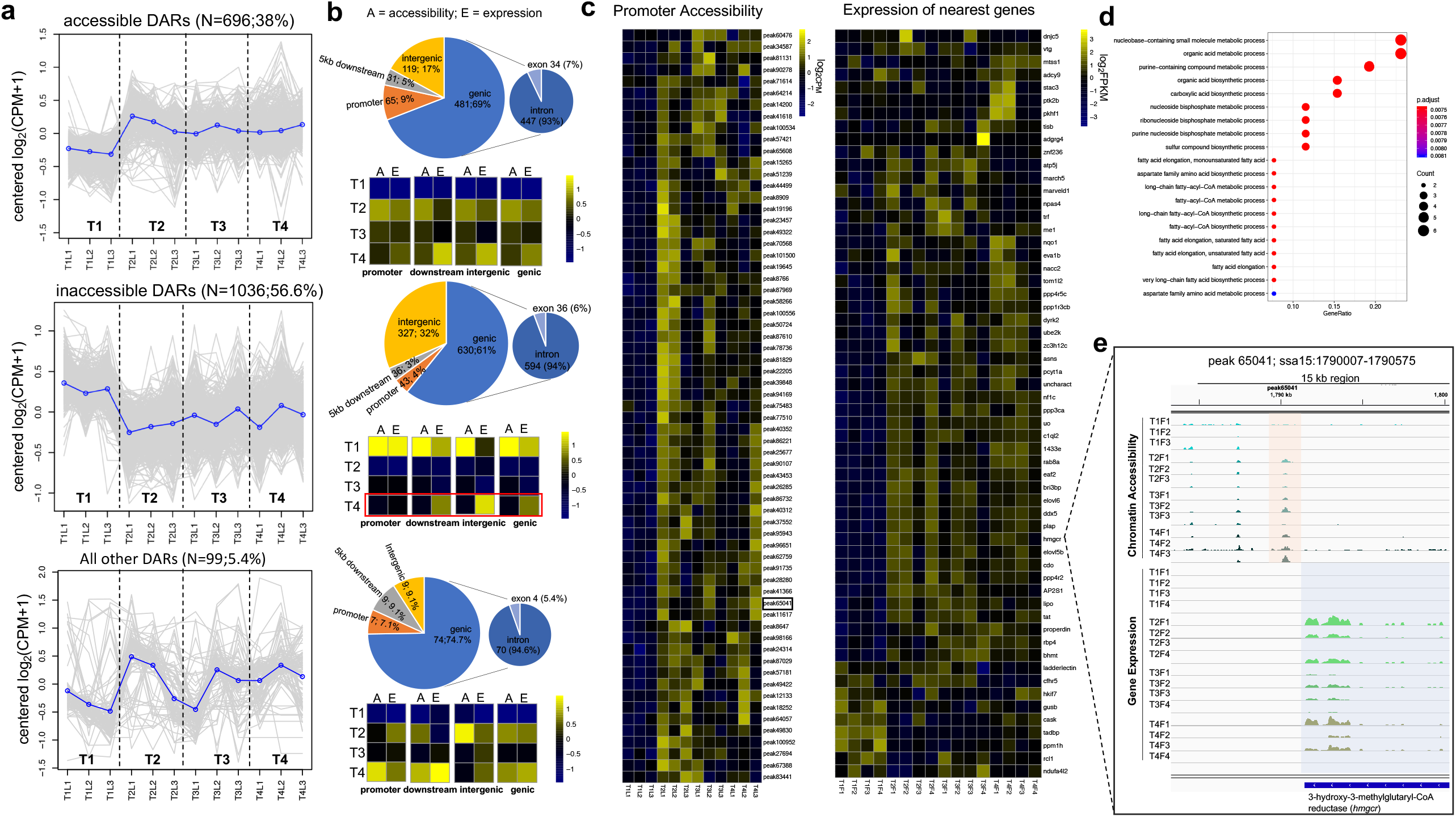
Dynamics of chromatin accessibility and gene expression as a function of genomic location reveals strong association with putative cis-regulatory elements (CREs). **a**, co-accessibility analysis reveals stable chromatin states following the onset of the maturation. The majority of DARs exhibited either reduced (n=1036 or 57%) or increased (n=696 or 38%) accessibility at T2 and remained unchanged at later timepoints. Less than 10% of DARs (n=99) displayed an oscillating pattern. The y-axis in each graph represents the mean-centered log2(CPM+1) value across time points on the x-axis. Accessibility of single DAR is plotted in grey, while the mean accessibility of each cluster is plotted in blue. **b**, Genomic distribution of DARs across gene models in the salmon genome, promoter defined as genomic regions located 5kb upstream of TSSs and multi-omic heatmaps showing mean accessibility and expression at each of the four time points per genomic location. **c**, Heatmaps 65 DARs located 5kb upstream of TSSs representing cis-regulatory elements (CREs) that exhibit increased accessibility at T2 along with gene expression of their target (nearest) genes. This showed upregulation of the majority of the associated genes in a tightly coordinated manner (n = 46; 79%’ *Χ*^*2*^ *p* < 8.028^-08^). **d**, Enriched gene ontology (GO) terms (hypergeometric test, Bonferroni-adjusted *P* < 0.05) among the list of the CREs-regulated genes. This list includes genes involved in hepatic lipid metabolism (*hmgcr*) and energy metabolism (*elovl5b* and *elovl6*). **e**, IGV visualisation of a 15 kb region of Ssa15 spanning the CRE and exons of the *hmgcr* gene provides fine-scale view of the coordinated gene expression response to increased accessibility.

### Differential chromatin accessibility strongly correlates with bidirectional regulation of global gene expression

To begin exploring the relationship between chromatin accessibility changes and gene expression, we first mapped the genomic location of DARs and found the majority were located in genes (65%) or within 5 kb upstream (6%) or 5 kb downstream (4%), affirming the quality of the ATAC-seq dataset. We also detected a quarter of DARs (25%) located on average 36 kb distal to their nearest gene, a low proportion in comparison to domesticated terrestrial livestock ^40^. Starting with the subset of DARs located in genes (exons and introns) we found the proportion of variation in gene expression explained by chromatin accessibility changes was high (Fig. 4c, d). For example, chromatin state changes at T2, compared with T1, explained 56% of the variation in gene expression using linear regression. The dynamic was bidirectional, with accessibility changes associated with both up and down regulation of global gene expression, and strongest at the early timepoint T2 (Fig. 4c). We repeated the analysis for DARs located within 5 kb of transcription start sites to assess the strength of association with physically proximal putative cis-regulatory elements (CREs). These had even higher association, explaining approximately 60% of the variation in global gene expression (Supplementary Figure S15). Together, this clearly demonstrated chromatin state changes played a dominant role in directing global changes in gene expression.

### Cis-regulatory elements regulate metabolism genes via chromatin state changes

To examine the biological consequence of chromatin state changes, we focused on CREs given their established role on transcriptional regulation via transcription factors (TFs) binding (41). We focussed on the subset of CREs that underwent a change in accessibility during the time course to evaluate i) the expression behaviour of their closet gene; ii) the biological function of those genes, and iii) any enrichment for transcription factor binding sites. We found a small subset of CREs (n = 65) underwent increased accessibility early in the time course and the majority (n = 46; 79%’ *Χ*^*2*^ *p* < 8.028^-08^) upregulated their nearest gene in a tightly coordinated manner (Fig. 5a – c). It also appears CREs more tightly controlled the downregulation of genes compared to DARs located in gene bodies, downstream regions or within intergenic regions (Fig. 5b, red box; Supplementary Table S16). The gene set associated with coordinated up regulation (Fig. 5c) exhibited significant GO enrichment related to lipid metabolism and energy metabolism (AAcyl-CoA biosynthesis) (Fig. 5d). Acyl-CoA are coenzymes involved in energy synthesis, consistent with the expectation of liver function through an energetically costly transition such as maturation.

To provide a fine-scale view of the coordinated response, a 15 kb region of Ssa15 spanning the CRE and exons of a gene that regulates hepatic lipid metabolism (*hmgcr*) ^42^ is provided in Figure 5e. The final analysis used HOMER to search for TF motifs that bind master regulators driving gene transcription ^43^. Specifically, we computed the enrichment of TF motifs in CREs that gained chromatin accessibility against a background that remained inaccessible. This revealed significant enrichment of 13 motifs, corresponding to the preferred binding sites of specific transcription factors present in 29 - 58% of targets following the onset of maturation (Supplementary Figure S16). Among these, the most significant motif matched the global transcriptional regulator E3 Ubiquitin-Protein Ligase CNOT4 that regulates essentially every aspect of gene expression, from mRNA synthesis to protein destruction including the degradation of RNAPII ^44^. The results strongly suggest that chromatin state changes at CREs directly control gene expression in liver and upregulate energy metabolism genes via changes in TF activity.

### Multiomics data integration using gene regulatory networks

The final component of our analysis sought to co-analyse all available data to infer gene regulatory networks (GRNs) responsible for the onset of maturation. GRNs provide a platform for integrating multiomic data and can be used to characterize the dynamics of perturbations during biological transitions such as puberty and other complex traits ^45,46,47,48,49^. Here, we used the approach to co-analyse genes with evidence of differential behaviour using seven categories that included expression (DEGs), changed methylation at gene bodies (DMGs) or promotors (DMPs) and differential chromatin accessibility (DACs). To focus the analysis towards investigation of key regulators, we also performed regulatory impact factor (RIF) analysis. This used co-expression correlation between TFs and their target differentially expressed genes to identify 305 significant regulators (Supplementary Table S17; Supplementary Figure S17). Of the seven categories, the majority of 1,858 genes prioritised for GRN construction were DEG (n = 1,400) or DMG (n = 700). The overlap between categories, for example where genes were both DEG and DMG (n = 442), is given in Fig 6.a. The gene set showed significant GO enrichment (1 GO-CC and 8 GO-MF terms) to hormone activity and steroid hormone receptor activity (Fig. 6b) and their expression patterns showed clear tissue-specific clustering (Supplementary Fig.S18a) suggesting biological relevance to the trait. GRN construction using the 1,858 genes yielded 835,084 connections with a mean of 449 connections per gene. For visualisation, we only considered genes with significant correlations ≥ ±0.95 (929 gene with 17,708 connections) (Supplementary Fig.S18b). Most network genes (N=777, ∼42 %) belonged to pituitary compared to 33% and 25% in ovary and liver. These figures also were reflected in the number of connections per tissue (Supplementary Fig.S18c). Genes with the highest change in the number of connections are likely to be key regulators, and the top 20 included five zinc finger proteins (Supplementary Table S18.) Two of these transcription factors, znf664 and znf239, were expressed in pituitary suggesting their key role in maturation onset. Interestingly, the most highly connected genes also included two uncharacterised (dark) Atlantic salmon genes (106590493, 106612553) that displayed maximum expression in ovary.

**Fig.6.**
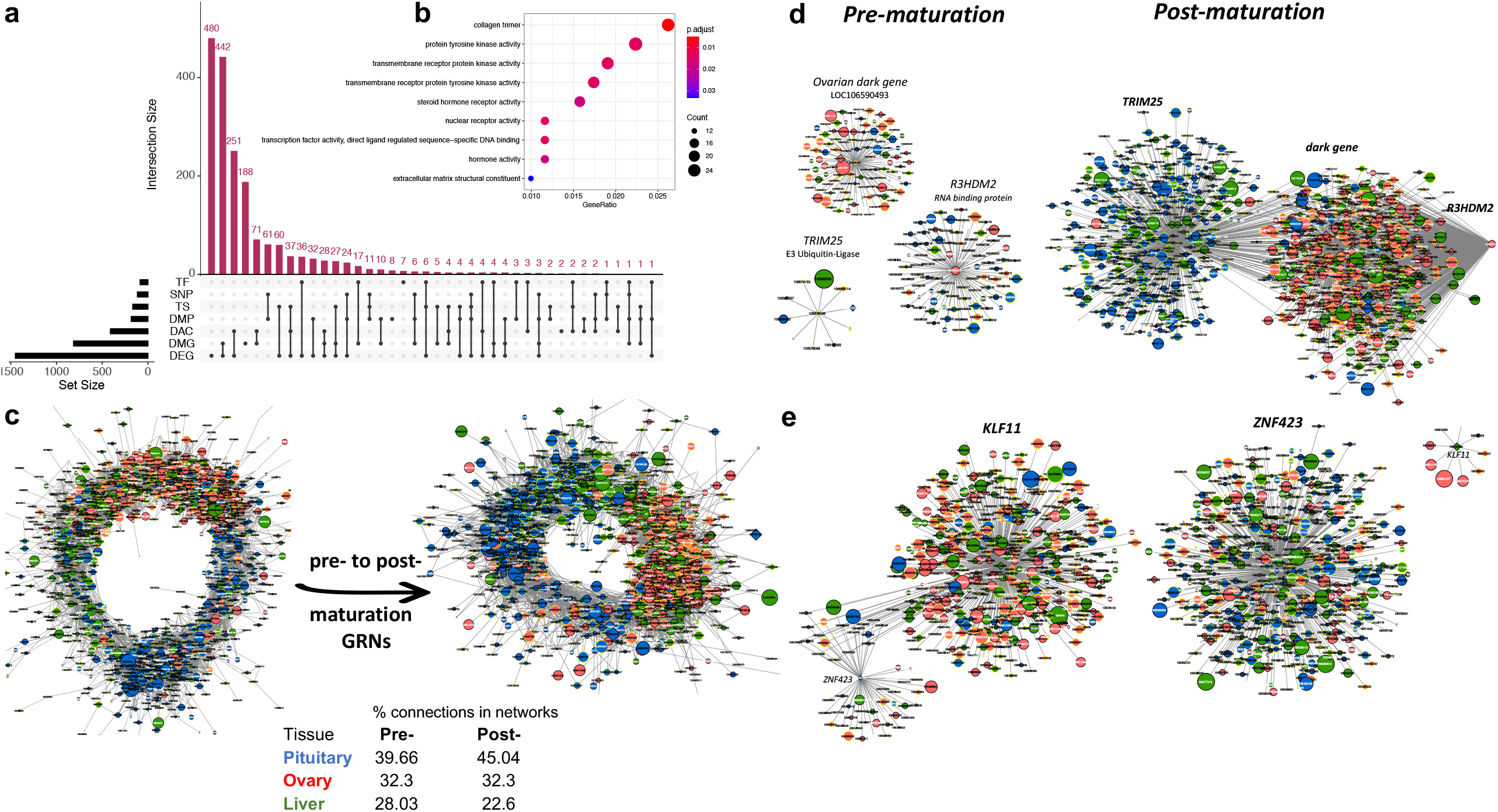
Gene regulatory networks (GRNs) constructed via integrating multiomics results. **a, UpSet plot showing the intersection among the 1,858 genes selected for network analyses.** Genes were selected when represented at least once and have a mean normalised expression of at least 0.2 FPKM. Network genes were divided into 7 categories: differentially expressed genes (DEG), differentially expressed genes/promoters (DMG, DMP), differentially accessible regions (genic and promoter regions as DAC), genes harbouring SNPs reported in literature as being associated with salmon maturation (SNP), tissue-specific genes (TS) and key regulators identified by RIF (TF). The nature of a given intersection is indicated by the dots below the bar plot. For example, the 442 genes in the second column are both differentially methylated and differentially expressed but not found in other categories. **b, Enriched gene ontology (GO) terms** (hypergeometric test, Bonferroni-adjusted *P* < 0.05) among the list of 1,858 network genes along with the gene ratio for the genes that map to each term. The majority of the enriched terms are related to hormone and receptor activities. **c, GRNs constructed using the PCIT algorithm for the pre- and post-maturation samples**. For visualisation purpose, only the most significant 10% of correlations their respective genes were considered. This shows increases in the connections in the pituitary that constituted 45% of the post-maturation network connections. All nodes are represented by ellipses except for genes coding key regulators (TFs) have diamond shape. Nodes with yellow borders are differentially methylated, whereas nodes with white labels are differentially accessible between pre- and post-maturation samples. Node colours are relative to the tissue of maximum expression with blue represents the pituitary, red represents ovary and green represents liver. The size of the nodes is relative to the normalized mean expression values in all samples. **d, Subnetworks of top differentially connected genes**, the networks created with the most trio genes differentially connected between pre- and post-maturation networks. This revealed TRIM25, a E3 Ubiquitin ligase as the key regulators with the greatest number of gained connections in the post-maturation network. TRIM25 was highly expressed in the pituitary and underwent changes in DNA methylation. **e, Subnetworks of top differentially connected TFs**, networks created with the most differentially connected TFs between pre- and post-maturation networks showed zinc finger protein 423 (ZNF423) as the key regulator with the greatest number of gained connections and Kruppel-like factor 11 (KLF11) as the regulator with the least number of gained connections going from pre-to post-maturation.

### Differential GRN connectivity identifies key regulatory factors

Key regulators are likely to undergo substantial change in their number of connections and identify gene networks driving the transition to maturation. This prompted construction of separate networks using pre- and post-maturation stage data, before identifying those genes that underwent the largest change in connectivity (pipeline workflow is provided in Supplementary Figure S1b). For visualisation purposes, we only included 10% of the most significant connections that included 1,412 genes with 17,260 connections in the pre-maturation GRN and 1,310 genes with 22,059 connections in the post-maturation GRN (Fig. 6c). Next, we computed the differences in the patterns among the tissues comprising the two networks (Fig.6c). The pituitary gland and ovary had the most abundance (∼ 45% and 32%, respectively) of connections compared to a lower percentage of connections (∼23%) in liver after maturation. We computed the differential connectivity for all genes and identified the most differentially connected genes (DCGs) (n=186 genes; 10%) (Supplementary Table S18) between the pre and post-maturation networks with more connections: 80,536 in post-maturation compared to pre-maturation network with 56,971. These were mainly expressed in pituitary (44%) and most connections involved DEGs (74%) and DMGs (47%). Finally, we identified regulators that gained the most connections post-maturation (Table 1, Fig. 6). The top ranked regulator was *trim25* (encodes a ubiquitin E3 ligase) and underwent a profound change in connectivity (from 10 to 599). *znf423* is a ZF-TF with multiple roles in signal transduction during development. It was predominantly expressed in pituitary and contributed to 3 categories in the network (DEG, TF, DAC) with multiple roles in signal transduction during development and a salmon dark gene highly expressed in ovary (Fig. 6d). Then, we focused on TFs contained among the top 10 regulators that were differential connected genes between the pre and post-maturation networks. This revealed *znf423* to be the most differentially connected TF (from 58 to 576) and Krueppel-like factor 11 (klf11) as the least differentially connected TF (from 513 to 9) (Fig. 6e; Table 1). Enrichment of motifs for a ubiquitin E3 ligase (*cnot4*) and ZNF TF (Zic) among ATAC-seq signals at CREs confirms their regulatory roles. Several studies have previously demonstrated roles for ZNF factors in controlling onset of female puberty in many species including humans ^50,51,52,53^.

**Table 1.**
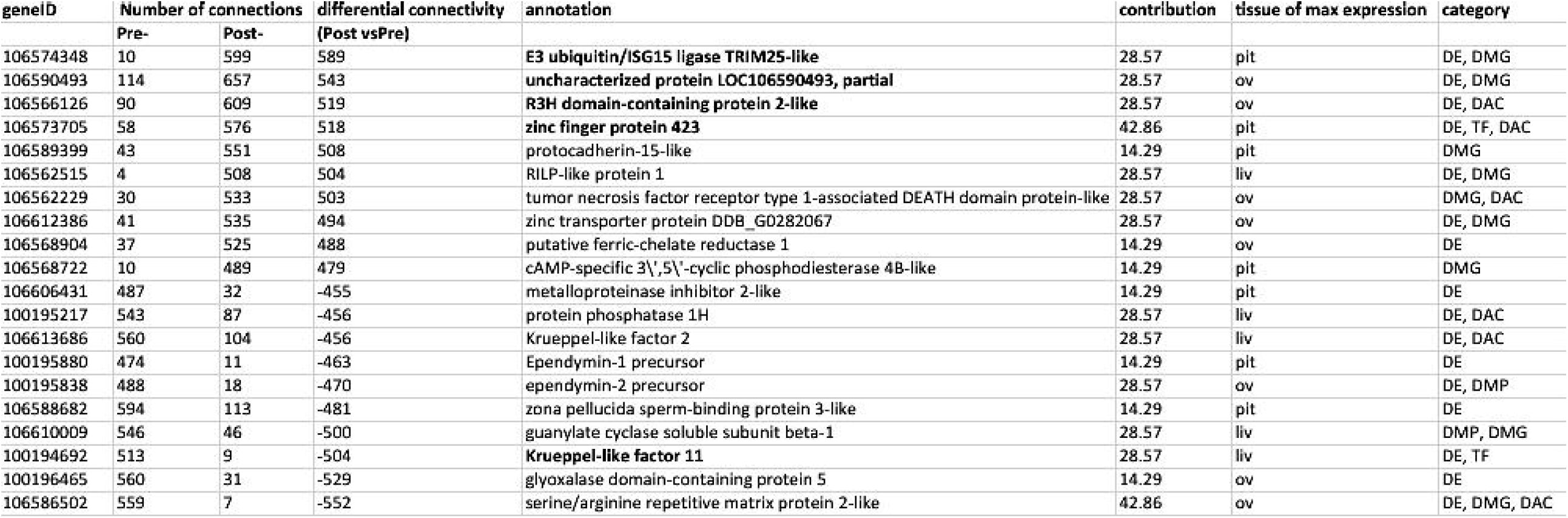

## Discussion

Identifying the biological mechanisms controlling complex traits is a sizable challenge. We designed our study on the assumption that the dynamic network of molecules coordinating the spatiotemporal changes driving sexual maturation would be inaccessible to investigation using only a single layer of “omics”. We anchored the study around the collection of tissue transcriptomes to visualize their changing circuitry across the period where Atlantic salmon commence their trajectory towards sexual maturation. Importantly, we also characterized the changing epigenomic landscape through interrogation of DNA methylation and chromatin state changes. Integration of the resulting multiomic dataset used rigorous quantitative approaches, and when performed inside the context of a defined biological transition, has given us an unprecedented ability to characterise the onset of maturation at the molecular level in a non-model species of worldwide aquaculture and ecological importance.

The availability of dimensional data allowed us to identify the dominant epigenomic changes controlling gene expression. We conclude that global changes in DNA methylation had little predictive power to explain changing gene expression beyond a small subset involved in chromatin remodelling. While methylation changes are a striking feature of embryonic development, they appear not to have been responsible for the rapid and numerous changes in gene expression documented here. Conversely, we observed high correlation between chromatin state changes and altered gene expression for the single tissue with ATAC-seq data (liver). The correlation was highest for differentially accessible regions immediately adjacent to coding genes, implicating *cis*-regulatory elements.

The identification of key genes relied on characterisation of differential behaviour using samples collected before, and after, fish were subjected to a photomanipulation trigger designed to stimulate maturation. Multiple data patterns confirm we successfully initiated early stage maturation. Increasing average gonadosomatic index demonstrates a physiological response occurred, and this was paralleled by significant global upregulation of gene expression in the ovary and a more modest remodelling of the pituitary transcriptome. Together, this provided confidence that the characterised DEG, DMG and DAC patterns are likely to successfully implicate genes directly involved in maturation. We showed upregulation of pituitary hormones including gonadotropins along with other pituitary genes involved in a range of reproduction related functions including steroidogenesis. Differentially methylated genes were enriched for follicular development and the control of gonadotropin-releasing hormone excitability. Integrated transcriptome and methylome analysis in ovary implicated chromatin remodelling genes in controlling maturation. Finally, differentially accessible CREs in liver were enriched for lipid metabolism and energy metabolism genes.

Despite the advantages of the multiomic approach used, limitations may be imposed by the range of tissues, timepoints and technical features of the assays used. For example, we treated tissues as homogenous entities in an approach that ignores the spectrum of constituent cell types and their differentiated roles that single-cell multiomic studies have begun to explore ^17,54^. Further, suboptimal partitioning of the brain at dissection hampered our ability to assign the role of the hypothalamus separately from the brain stem, cerebellum and olfactory bulb. Consequently, the variation in whole brain transcriptomes among replicates within timepoint was so large as to prevent meaningful analysis. Finally, we were unable to generate high quality chromatin state data from ovary samples despite repeated attempts. An incomplete compendium of tissues and datatypes has resulted in an imperfect view of the changing epigenomic landscape.

The promise of multiomic data will remain unfulfilled without methodological approaches capable of identifying system perturbations associated with phenotypic change. Here, we used a gene regulatory network approach which has proven successful for investigation of puberty and other complex traits ^47,48,49^. We identified t*rim25*, a gene encoding a ubiquitin E3 ligase, as the network element undergoing the most dramatic change in connectivity and strongly suggests it plays a key role. Looking forwards, it is tempting to speculate that a number of the genes identified represent high value targets for manipulation via gene editing in an attempt to delay or ablate sexual maturation. Among the range of putative targets identified, the core components of the SWI/SNF chromatin remodelling complex (*arid1B* and *smarca2*) are appealing due to their ability to exert wide ranging change in gene expression. The results described may therefore lead to better management of unwanted early maturation within an aquaculture setting where the completion of maturation is associated with reduced product quality and production inefficiencies.

## Supporting information

STable1

STable2

STable3

STable4

STable5

STable6

STable7

STable8

STable9

STable10

STable11

STable12

STable13

STable14

STable15

STable16

STable17

STable18

STable19

## Methods

### Induction of maturation through photoperiod manipulation and tissue sampling

Animals were managed using photoperiod manipulation to synchronise the timing of commitment into maturation. A population of female brood stock were used that were ∼ 36 months post fertilization in April 2017. The management of the animals and associated timeline for sampling events is given in Fig.1a. In order to measure and control for variation between individuals, 4 fish (biological replicates) at each of the four time points (T1-T4) were used. The maturation status of animals (leading up to the long day photoperiod initiation) was monitored by ultrasound. Control samples at T1 time point were collected on mid-June 2017 before induction of maturation occurred late-June 2017. Following the application of the long photoperiod, tissues were sampled at different three time points in 2 weeks intervals (T2-T4). At each sampling event, the gonadosomatic index GSI was calculated from the ovary mass as a proportion of the total body mass as follows: GSI = [ovary weight / total body weight] × 100.

### RNA isolation, RNA-seq library preparation and sequencing

Tissue samples were preserved in RNA-Later at −80 °C and total RNA was isolated using RNeasy mini kit (QIAGEN) as previously described ^55^. Tissues were lysed twice in 450 µL of lysis solution on a Precellys 24 homogenizer for 30s at 4.0 ms^−1^. RNA was bound to a column and washed twice before elution with 40 µL at room temperature. RNA quantity and quality were assessed using a NanoDrop ND-1000 spectrometer, Qubit 2.0 fluorometer and Agilent 2100 bioanalyzer. Messenger RNA (mRNA) was isolated from 1 µg of total RNA. 64 RNA-Seq libraries (4 time points x 4 tissues x 4 biological replicates) were prepared using the TruSeq RNA Sample Preparation Kit (Illumina). Libraries were sequenced on Illumina Nova-Seq 6000 sequencing platform at the Australian Genome Research Facility (AGRF) in Melbourne, Australia. Sequencing produced a total of 4.4 billion individual 150 bp paired-end reads and ∼ 70 million PE reads per library (Supplementary Table S1).

### Transcriptomic data quality control (QC), genome mapping and read counting

Illumina reads were checked for quality using FastQC software. High quality reads (Q>30) were mapped to the Atlantic salmon genome ICSASG_v2 ^56^ using TopHat2 version 2.1.1 ^57^ with default parameters. Alignment files in BAM format were sorted by read name and converted into SAM format using SAMtools version 1.4 ^58^. The Python package HTSeq version 0.7.2 ^59^ was applied to count unique reads mapped to exons using default parameters except for “*reverse*” with the strandedness.

### Differential gene expression and clustering analyses

Raw counts were analysed using the edgeR package ^60^ in the R statistical computing environment to infer differential gene expression among tissues. The four tissues at the long photoperiod time points (T2, T3 and T4) were compared to the control samples at T1. *P*-values for differential gene expression were corrected for multiple testing using the Benjamini and Hochberg algorithm ^61^. For further analyses of differential expression, only genes with a false discovery rate (FDR) of < 0.05 and have at least absolute log2(fold change) > 1 were considered significant. PCA was conducted on the lists of significant DEGs using normalised expression data (log2FPKM) using the function *--prin_comp* within trinity. Hierarchical clustering analysis was conducted using trinity’s utility *analyze_diff_expr*.*pl* on significant DEGs in each tissue where mean-centred normalized expression (log2-transformed FPKM+1) were compared across time points ^62^. Gene clusters with similar expression patterns were obtained using the Perl script *define_clusters_by_cutting_tree*.*pl* within trinity to cut the hierarchically clustered gene tree into clusters with similar expression using the *--Ptree* option.

### Gene Ontology (GO) enrichment of the identified gene clusters

To infer the functions of the gene clusters, gene ontology (GO) enrichment was performed to identify the enriched biological themes using the R package clusterProfiler version 3.9 using default settings ^63^. The ENTREZ gene identifiers of up- and downregulated clusters per tissue were used as query gene list against the background genes in each tissue. For the purpose of the enrichment analysis, GO categories with a corrected *P*-value of < 0.05 were considered significant. Categories of candidate genes implicated in maturation were visualised as heatmaps using their normalised expression values with the R package pheatmap version 1.0.12 https://cran.r-project.org/web/packages/pheatmap/pheatmap.pdf.

### Genomic DNA isolation, WGBS library preparation and sequencing

Tissue samples were snap frozen in liquid Nitrogen and stored at −80 °C until genomic DNA (gDNA) was extracted using DNeasy blood and tissue kit (QIAGEN). Tissues were lysed in 360 µL of lysis solution on a Precellys 24 homogenizer for 30s at 4.0 ms^−1^. Samples were incubated with 40 µL of Proteinase K enzyme at 56 °C for 1 h. Following lysis, samples were treated with RNase (8 µL of RNase A incubated for 2 min at room temperature). DNA was bound to the provided columns, washed twice and eluted in 100 µL at room temperature. gDNA purity were assessed by gel electrophoresis and NanoDrop ND-1000 spectrometer. DNA concentration and integrity were assessed using Agilent 2100 bioanalyzer. gDNA was fragmented (200-400bp) by sonication using Covaris S220, followed by end repair/adenylation and adapter ligation. Bisulfite modification was performed to the DNA fragments using the EZ DNA Methylation-GoldTM Kit (Zymo Research, Inc.). Twelve libraries prepared from 3 tissues (pituitary, ovary and liver), 2 time points (T1 and T4) and 2 biological replicates. Libraries were sequenced on HiSeq 2500 sequencing platform at Novogene, Hong Kong. Sequencing produced a total of 2.2 billion individual 150 bp paired-end reads and 185 M PE reads per library. Bisulfite conversion rates (percentage of C changed to T after bisulfite treatment) were consistently >99.8%.

### WGBS data QC, genome mapping and methylation calling

Raw data quality control was performed using Trim Galore v0.5 (http://www.bioinformatics.babraham.ac.uk/projects/trim_galore/) to filter bases (Q scores < 30) and remove both universal and indexed adapter sequences. Processed high-quality data were mapped to into a bisulfite-converted version of the Atlantic salmon reference genome ICSASG_v2 ^58^ using BSseeker2 v2.1.8 ^66^ with default parameters for aligning paired-end libraries using Bowtie2 (Langmead & Salzberg 2012). PCR duplicates were detected and removed using Picard MarkDuplicates (http://broadinstitute.github.io/picard/). Filtered (duplicates-free) reads (110 M PE reads) were retained for downstream methylation analysis with an average genome coverage of 11x in pituitary, ovary and liver. Methylation calling was conducted using the Python script *call-methylation*.*py* within BSseeker2. CGmap files were used for subsequent exploratory and differential methylation analyses. The *mstat* command within CGmap tools was used to generate global and CG context (CG, CHG, CHH) DNA methylation levels ^66^.

### DNA methylome exploratory analyses

As CG methylation contributed to the bulk of methylated Cs, average methylation levels of genome-wide CpG positions were calculated in 50 kb bins across the genome using *mbin* command within CGmap tools and plotted as Violin plots using the R package vioplot, https://cran.r-project.org/web/packages/vioplot/index.html. To begin assessment of the quality of our libraries, common CpGs with minimum 10x coverage among the 12 samples were used in PCA using *prcomp* implemented in R. Correlation matrices (based on Pearson coefficient) were prepared using the R package *corrplot* (https://cran.r-project.org/web/packages/corrplot/index.html). Hierarchical clustering analysis was conducted with *hclust* implemented in R using compute linkage and Euclidean distances.

### Differential CpG methylation analysis

The R package DSS was used to identify differential methylation regions using common CpGs ^67^. In each tissue, two replicates at T4 were compared to the control samples at T1 based on CpG methylation levels. At each CpG site, the methylation (M) level was calculated as a proportion of the total counts (coverage) as follows: M levels = [methylated counts / total counts] × 100. DSS was selected as it takes into account the biological variation among replicates (characterized by a dispersion parameter) and the sequencing depth. Differentially methylated loci (DMLs) were identified by estimating mean methylation for all CpG sites followed by estimating dispersion at each site and conducting a Wald test (P < 0.001). Smoothing (combining information from nearby CpG sites to improve the estimation of methylation levels) was utilised to obtain mean methylation estimates in WGBS data where the CpG sites are dense. Based on the DML results, regions with statistically significant CpG sites were identified as a differentially methylated regions (DMRs) with minimum length/distance of 50 bp and minimum CpG coverage of 3. Mean methylation between groups of greater than 10 % (delta = 0.1) and *P* < 0.001 was considered significant. A circos plot was produced to visualize multi-tissue genome-wide DMRs using Circos (http://circos.ca/software/). Individual DMRs were also visualized using the *showOneDMR* function within the DSS package to plot both the methylation percentages (including a smoothed curve) as well as the coverage depths at each CpG site.

### DMR annotation, DMGs and DEGs correspondence analysis

Differentially methylated regions were compared against the protein coding gene set annotated on reference ICSASG_v2 using custom Perl scripts. This classified DMRs as overlapping a gene body (genic), 5kb upstream of a transcription start site TSS (putative promoter), 5 kb downstream of TSS (5kb downstream), or otherwise intergenic. The distance between each DMR and nearest gene is provided in Supplementary tables 9 – 11. The overlap between significant genes from differential expression and methylation was checked using the *intersect* function within bedtools ^68^.

### GO enrichment of DMGs

GO enrichment analyses were conducted on both the sets of hypermethylated genes (n= 1,156) and genes found to be hypermethylated and upregulated in ovary (n= 148) using the R package clusterProfiler. Genes driving GO enrichment were plotted as a heatmap using the R package pheatmap as above.

### Nuclei extraction, ATAC-seq library preparation and sequencing

ATAC-seq libraries were prepared from frozen tissues using the Omni-ATAC method ^69^ with the following modifications. Frozen tissue (20 mg) was ground in liquid nitrogen using a mortar and pestle. The pulverized tissue was transferred to a pre-chilled 2 ml dounce homogenizer containing 1mL cold 1x homogenisation buffer and homogenised with the pestle to form a uniform suspension (10-20 strokes). The homogenate was filtered with a 40uM nylon cell strainer (BD Falcon) before layering onto the iodixanol solution as described previously ^69^. The ratio of nuclei to enzyme concentration was optimised for each sample by performing transposition reactions containing 50000, 100000 and 200000 nuclei with 2.5ul of tagment enzyme in 50ul of transposition mix ^69^. The transposed DNA was amplified with custom primers as described elsewhere ^70^. before libraries were purified using Agencourt AMPure XP beads (Beckman Coulter) and quality controlled using a Bioanalyser High Sensitivity DNA Analysis kit (Agilent). Twelve liver ATAC-seq libraries arising from 3 biological replicates x 4 time points (T1-T4) were sequenced at the IMB sequencing facility (University of Queensland) on an Illumina NextSeq 150 cycle (2 × 75 bp).

### Chromatin accessibility data QC, genome mapping and peak calling

Sequencing produced a total of 1.2 billion individual paired-end reads (Supplementary Table 14). Raw reads were mapped to the Atlantic salmon reference genome ICSASG_v2 ^58^ using BOWTIE2 version 2.3.5.1 with the *--very-sensitive* parameter ^71^. Duplicate reads were removed using the MarkDuplicates function in Picard (http://broadinstitute.github.io/picard/). Multi-mapped reads and mitochondrial reads were filtered out and only uniquely mapped reads (MAPQ > 10) were extracted from alignment files using SAMTOOLS for downstream analyses.

For peak calling, the model-based analysis of ChIP-seq (MACS2) (https://github.com/macs3-project/MACS) was used to identify read enrichment regions “peaks” using default parameters. Only peaks detected in at least two replicates per condition were used for downstream analyses, and peaks across timepoints were merged to generate a unique peak list per tissue. The number of raw reads mapped to each peak was quantified using the Python package HTSeq version 0.11.1 ^59^.

### Differential accessibility and clustering analyses

Samples from the long photoperiod time points (T2, T3 and T4) were compared to control samples (T1) for each tissue. Raw counts were analysed using the R package edgeR and P-values were corrected for multiple testing using the Benjamini and Hochberg algorithm. Peaks with FDR < 0.05 and log2FC > ± 1 were considered significantly differentially accessible regions (DARs). PCA of significant DARs used normalised accessibility data (log2CPM) prepared using the function *--prin_comp* within trinity. Hierarchical clustering analysis was conducted using *analyze_diff_expr*.*pl* where mean-centred normalized accessibility (log2CPM+1) were compared across time points ^62^. Gene clusters with similar accessibility patterns were obtained using the Perl script *define_clusters_by_cutting_tree*.*pl* to cut the hierarchically clustered gene tree into clusters with similar accessibility patterns as described above.

### Genomic distribution of DARs within clusters

Hierarchical clustering identified both accessible and inaccessible DAR clusters. DARs per cluster were annotated in a genomic context (genic, promoter, 5 kb downstream or intergenic) as previously done for annotation of DMRs.

### ATAC-seq and RNA-seq correspondence analysis

Only DARs co-located with genes and promoters were used for co-analysis with gene expression data. The relationship between accessibility of DARs and gene expression was visualised by overlying information of significant DARs to genome-wide normalised expression estimates in liver samples and plotted as a MA-biplot. A linear regression analyses were performed to assess correlations between accessibility and expression abundance and the effect of changes in accessibility and changes in gene expression across time. Chromatin accessibility and gene expression data were visualised using Gnuplot version 5.0.7 (http://www.gnuplot.info) by overlying accessibility data of significant DARs at genes and promoters to genome-wide normalised expression estimates at each timepoint.

### Multiomic heatmap analysis per time and genomic regions

All heatmaps were produced using the R package pheatmap. GO enrichment analyses have been conducted on the set of nearest genes to accessible promoters using the R package clusterProfiler as described above. The integrated genome viewer (IGV) was used to visualise the relationship between accessibility and gene expression in a 15kb region that contains *hmgrc* gene and its promoter region.

### Motif enrichment analyses

The function *findMotifsGenome*.*pl* within Homer software version 4.11 (http://homer.ucsd.edu/homer/) was used with default parameters to find sequence motifs significantly enriched among accessible DARs vs inaccessible DARs located within promoter regions.TF motifs that are highly enriched (*P* value < 1 × 10^−10^) were selected.

### Inference of master regulators

Master regulator analysis was performed using regulatory impact factor (RIF) metrics described by ^72^ to identify key regulators contributing to the differential expression in the T4 vs T1 comparison in each tissue. Data for potential transcription factors (TFs) in Atlantic salmon were taken from Mohamed et al., 2018. As most of the transcriptional changes were detected at T4, RIF was applied to the T4-T1 comparisons for each tissue. Briefly, RIF exploits the differential co-expression concept where regulators were contrasted against unique lists of genes that were differentially expressed at T4 in each tissue. Genes with a mean expression FPKM < 0.2 were excluded. Those scores deviating ± 2.57 standard deviation from the mean were considered significant at *P* < 0.01. We identified a total of 305 significant regulators (113, 68 and 123 in pituitary, ovary and liver, respectively at *P* > 0.01). Most of these regulators (n=298; 97.7%) were unique to each tissue leaving only 7 that were shared among tissue pairs (Supplementary Table S17; Supplementary Figure S17). The regulators identified were used as input for construction of gene regulatory networks as summarized in Supplementary Figure S1b.

### Gene regulatory network (GRN) analysis

Genes from different omics analyses (DEGs, DMGs, DMPs, DACs) along with key transcription factors identified by RIF (TFs), as well as information for tissue-specific (TS) genes and gene-harbouring GWAS SNPs (SNPs) were selected based on overlap (at least once) and mean normalised expression (at least 0.2 FPKM) for network construction. The R package UpSetR (https://cran.r-project.org/web/packages/UpSetR/vignettes/basic.usage.html) was used to investigate the cross-talk among genes from different sources. For gene network inference, genes were used as nodes and significant connections (edges) between them were identified using the Partial Correlation and Information Theory (PCIT) algorithm ^73^, considering all samples. PCIT determinates the significance of the correlation between two nodes after accounting for all the other nodes in the network. Connections between gene nodes were accepted when the partial correlation was greater than two standard deviations from the mean (*P* < 0.05). The output of PCIT was visualized using Cytoscape Version 3.7.2 ^74^.

In order to explore differential connectivity during maturation onset, two networks were created; one using 12 samples at T1 (pre-maturation) and a second using 36 samples at T2, T3 and T4 (post-maturation). The number of connections of each gene in each network was computed, making it possible to compare the same gene in the two networks to identify differentially connected genes (DCGs). From these networks, we explored a series of subnetworks. First subnetworks based on the top trio genes and the top regulators (TFs) based on their differential connectivity between pre-and post-maturation. Pre- and post-maturation networks were constructed from the 12 control samples at T1 and 36 post-maturation (T2-T4) samples.

## Data availability

Sequencing reads, raw and processed data used in this multiomic study have been submitted to the NCBI Gene Expression Omnibus (GEO) database under Accession numbers GSE157003 (combined data types); GSE157001 (RNA-seq); GSE156998 (ATAC-seq) and GSE156997 (whole genome bisulphite sequencing data). Gene network cystoscope file contains all networks described in the paper is supplied as supplementary file.

## Code availability

All codes used for gene expression (RNA-seq; https://github.com/AminRM/salmon_mat_transcriptomes), DNA methylation (WGBS;https://github.com/AminRM/DNAmethylomes), chromatin accessibility (ATAC-seq; https://github.com/AminRM/Salmon-Chromatin-Accessibility), gene networks (GRN; https://github.com/AminRM/maturation_GRN) analyses are available

## Acknowledgements

We acknowledge the cooperation and assistance provided by Saltas staff who performed the animal husbandry component of the experiment. The CSIRO Office of the Chief Executive provided financial support to Amin Mohamed. The work is dedicated to the memory of Harry King.

## Author Contribution Statement

AM, HK, BE and JK conceived and designed the experiment. AM generated and analysed RNA-seq and WGBS data. MM generated ATAC-seq data. AM and MN-S analysed ATAC-seq data. AM and AR analysed the gene regulatory networks. AM prepared all the figures/tables and prepared the first draft of the manuscript. All authors reviewed, commented on and approved the final manuscript.

## Ethics Statement

The fish were reared and euthanized in compliance with the CSIRO Animal Ethics Committee, under approval number 2017–02.

