## Supplemental information for "Integrated transcriptome, DNA methylome and chromatin state accessibility landscapes reveal regulators of Atlantic salmon maturation"

**Supplementary Results:**

**Salmon multi-tissue transcriptome profiling during maturation:**

A total of 4.4 billion 150 bp paired-end (PE) reads were obtained from 64 RNA-Seq libraries (~70 million PE reads per library). Hierarchical clustering of expression data was performed to examine the relationship among biological replicates within timepoints in each tissue. It also served to examine the relationship between samples across the time course, by comparing control samples prior to long photoperiod treatment (T1) with those following the onset of the long light treatment (T2, T3 and T4). Pairwise Spearman correlations revealed low variation among biological replicates and separation between samples from different timepoints for the ovary, liver and pituitary (**Supplementary Fig. 2**). Analysis of brain samples revealed much higher variability among biological replicates within timepoint, suggesting a low quality dataset that was excluded from subsequent analysis (**Supplementary Fig. 2**). We conclude the sampling of brain tissue was non uniform between fish, capturing different brain regions and generating the highly variable patterns in gene expression observed.

Transcriptome profiling of the pituitary revealed fewer differentially expressed genes (DEGs) compared with other tissues, however the number increased with increasing time following the onset of the long light regime. Only ten genes were differentially expressed at timepoint 2 (T2), increasing to 221 DEGs at T3 (75 and 146 genes were down- or upregulated) and 312 DEGs at T4 (76 and 236 genes were down- or upregulated). The range of $\log_2$(fold-change) and false discovery rates are summarized in

**Supplementary Fig. 2a** and **Supplementary Table 2**. The upregulated gene cluster showed significant GO enrichment with respect to 23 GO-BP (Biological Process), 13 GO-CC (Cellular Component) and 38 GO-MF (Molecular Function) terms related to maturation related functions as G protein−coupled receptor signalling and hormone activity (**Fig. 2c and Supplementary Table 5**). While 32 GO-BP and 10 GO-MF terms related to receptor ligand activity and developmental process involved in later stages of reproduction (derived from the presence of many genes of *zp3* encoding zona pellucida sperm-binding protein 3 that is essential for sperm binding during fertilization) showed significant enrichment among the downregulated gene cluster (**Supplementary Fig. 4; Supplementary Table 5**).

Transcriptome profiling of the ovary and liver employed the same analytical approach. This documented a radical transformation in gene expression occurred in the ovary, with differential expression of ~ 6000 genes at T2, T3 and T4 compared to the control T1 (adjusted $P < 0.05$) (**Fig 2a; Supplementary Fig. 3).** A total of 466 ovary genes were differentially expressed at T2 (63 and 403 genes were down- or upregulated). At T3, 1790 were differentially expressed (81 and 1709 genes were down- or upregulated). At T4, the ovarian transcriptome underwent extensive remodelling of 3737 genes (240 and 3497 genes were down- or upregulated) **(Supplementary Table 3)**. Hierarchical clustering of the ovary DEGs revealed distinctive expression profiles for post-maturation and identified two distinct clusters of upregulated genes (n=3476; 58% of ovary DEGs) and downregulated genes (n=301) (**Supplementary Fig. 3, 5a,b**).

The upregulated gene cluster showed significant GO enrichment with respect to 145 GO-BP, 15 GO-CC and 75 GO-MF terms related to cell adhesion, immune/inflammatory response, development (**Supplementary Fig. 5c;** Supplementary Table **6**). While 16 GO-MF terms related to channel activity were enriched among downregulated cluster (**Supplementary Fig. 5d;** Supplementary Table **6)**. The ovary DEGs are involved in several maturation-related functions such as steroidogenesis (*cyp17a1*, *sf-1*, *star*), hormonal receptors (*gnrhr2*, *amhr2*, *prlr*, *slr*), growth factors (*igf2*, *gdf11*, *tgfb3*), follicular development (*gata4*, *wnt5a*, *rspo1*, *foxl2*, *fstl1*, several sema and plexin genes), extracellular matrix remodelling (*fn1*, *cldn5* and several collagen chains) and immune/inflammatory response (*myd88t, tlr, cd2,igll1, ccl19, traf2*) (**Supplementary Fig. 6).**

In liver, 4001 genes were differentially expressed at T2 (1821, 2180 genes were down- and upregulated). At T3, 2153 were differentially expressed (957, 1196 genes were down- and upregulated). At T4, the liver transcriptome underwent extensive remodelling of 3387 genes (1301, 2086 genes were down- and upregulated) **(Supplementary Table 4)**. Hierarchical clustering of the liver DEGs revealed distinctive expression profiles for post-maturation and identified two distinct clusters of upregulated genes (n=3336) and downregulated genes (n=2347) (**Supplementary Fig. 7**). The upregulated gene cluster showed significant GO enrichment with respect to 45 GO-BP, 17 GO-CC and 23 GO-MF terms related to organic acid metabolic processes and mitochondrial transport (**Supplementary Fig. 7 and Supplementary Table 7**). While less terms were enriched among the downregulated cluster; 14 GO-BP, 13 GO-CC and 8 GO-MF terms related to translation and developmental process involved in reproduction mainly sperm-egg recognition during binding of sperm to zona pellucida (**Supplementary Fig. 7 and Supplementary Table 7**).

**Salmon multi-tissue DNA methylome profiling:**

Methylome sequencing produced 4.4 billion reads, with average coverage of ~18x for each of the 12 DNA methylome libraries (2 replicates x 2 conditions x 3 tissues). We found an average genome-wide methylation rate of 81% across all samples (**Supplementary Fig. 8; Supplementary Table 8)**, which is consistent with methylation range reported for other fish genomes (60-90%) (**Zhou et al., 2019; Ortega-Recalde et al., 2019**). We plotted the average CpG methylation levels in genomic bins to show the methylation levels, ranging from 0 to 1 (**Supplementary Fig. 8b)**. Clustering analysis of common CpGs in the 12 DNA methylome libraries revealed three main clades that grouped samples according to tissue type, but also separated the samples according to the maturation status (**Fig. 3a**). PCA analysis of common CpGs showed high variation among the ovary samples implying extensive remodelling in the ovarian methylomes (**Supplementary Fig. 8e**).

**Chromatin accessibility data:**

We obtained a total of ~750 million individual paired-end reads from 12 liver libraries (3 replicates x 4 time points). An average mapping rate of 93% was observed (duplicate free uniquely mapped reads) and 29 million qualified paired-end reads (sequencing fragments) per library were obtained. On average, ~94,000 high-confidence open chromatin regions (or peaks) were identified across liver libraries (**Supplementary Table S14**). These peaks were enriched around transcription start sites (TSSs) (**Supplementary Figure S13**).

#### Supplementary Tables

**Supplementary Table S1**: Description of multi-tissue transcriptome (RNA-seq) data. Sequencing raw data and mapping statistics for all samples at all time points.

**Supplementary Table S2**: Differentially expressed genes (DEGs) in pituitary (FDR < 0.05; logFC ±1) at T2, T3, T4 vs T1. Statistics for differential expression along with raw counts are provided for all pituitary samples.

**Supplementary Table S3**: Differentially expressed genes (DEGs) in ovary (FDR < 0.05; logFC ±1) at T2, T3, T4 vs T1. Statistics for differential expression along with raw counts are provided for all ovary samples.

**Supplementary Table S4**: Differentially expressed genes (DEGs) in liver (FDR < 0.05; logFC ±1) at T2, T3, T4 vs T1. Statistics for differential expression along with raw counts are provided for all liver samples.

**Supplementary Table S5**: Statistics associated with Gene ontology (GO) enrichment for each up- and down-regulated gene cluster in pituitary. Enriched GO categories BP, CC, MF were selected using a hypergeometric test at Bonferroni-adjusted P < 0.05.

**Supplementary Table S6**: Statistics associated with Gene ontology (GO) enrichment for each up- and down-regulated gene cluster in ovary. Enriched GO categories BP, CC, MF were selected using a hypergeometric test at Bonferroni-adjusted P < 0.05.

**Supplementary Table S7**: Statistics associated with Gene ontology (GO) enrichment for each up- and down-regulated gene cluster in liver. Enriched GO categories BP, CC, MF were selected using a hypergeometric test at Bonferroni-adjusted P < 0.05.

**Supplementary Table S8**: Description of methylome (WGBS) data, sequencing, mapping and methylation statistics.

**Supplementary Table S9**: Methylation statistics, genomic locations and annotation of pituitary differentially methylated regions (DMRs).

**Supplementary Table S10**: Methylation statistics, genomic locations and annotation of ovary differentially methylated regions (DMRs).

**Supplementary Table S11**: Methylation statistics, genomic locations and annotation of liver differentially methylated regions (DMRs).

**Supplementary Table S12**: Statistics associated with Gene ontology (GO) enrichment for hypermethylated genes in ovary. Enriched GO categories BP, CC, MF were selected using a hypergeometric test at Bonferroni-adjusted $P < 0.05$.

**Supplementary Table S13**: Statistics associated with Gene ontology (GO) enrichment for 148 (hypermethylated/upregulated) genes in ovary. Enriched GO categories BP, CC, MF were selected using a hypergeometric test at Bonferroni-adjusted $P < 0.05$.

**Supplementary Table S14**: Description of chromatin accessibility (ATAC-seq) data, sequencing, mapping statistics.

**Supplementary Table S15**: Differentially accessible regions (DARs) in liver ($FDR < 0.05$; $logFC \pm 1$) at T2, T3, T4 vs T1. Statistics for differential accessibility along with raw counts are provided for liver samples.

**Supplementary Table S16**: Genomic locations and annotation of liver differentially accessible regions (DARs) in accessible and inaccessible clusters.

**Supplementary Table S17:** Transcription factors identified as master regulators according to the regulatory impact factor (RIF) metrics in pituitary, ovary and liver tissues ($P < 0.01$).

**Supplementary Table S18**: Highly connected genes (Top 20) among the 1,858 genes selected for the gene regulatory network (GRN) construction. The table shows number of connections per gene, annotation, tissue of maximum expression and category.

**Supplementary Table S19**: Differentially connected genes (10%) between pre- and post-maturation GRNs. The table shows number of connections per gene (connectivity degree) in both pre-and post-maturation, differential connectivity (Post vs Pre), annotation, tissue of maximum expression and attributes for the different categories.

### Supplementary Figures

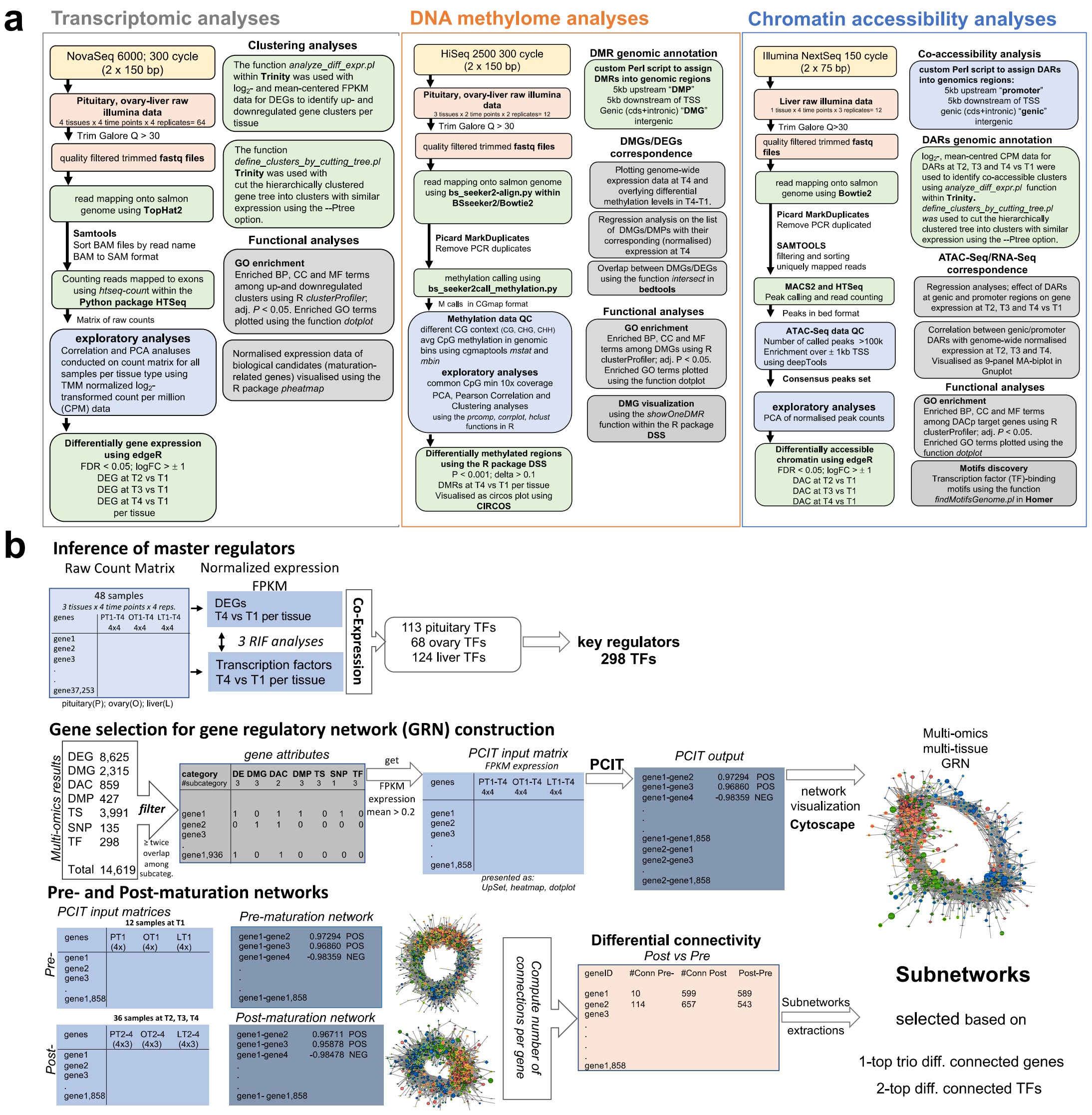


**Supplementary Figure S1.** Schematic view of the bioinformatic workflow for **a** Analyses of RNA-seq, WGBS and ATAC-seq data. Illumina reads were mapped onto the Atlantic salmon genome and alignment files were processed to extract gene expression counts, CpG methylation, peak counts for downstream analyses of differential analyses using different packages in the R environment. and **b** Inference of master regulators and gene regulatory network (GRN) construction. RIF metrics were used to identify master regulators contributing to differential expression observed at T4 in pituitary, ovary and liver. The unique list of these regulators were integrated with other multi-omics results that included differentially expressed genes from three tissues (DEGs), differentially methylated genes and promoters from three tissues. Differentially accessible chromatin (co-located with gene bodies and promoters) from liver. Along with data describing genes-harbouring SNPs and tissue-specific (TS) genes obtained from previous maturation GWASs and tissue-specific transcriptomes derived from the same samples.

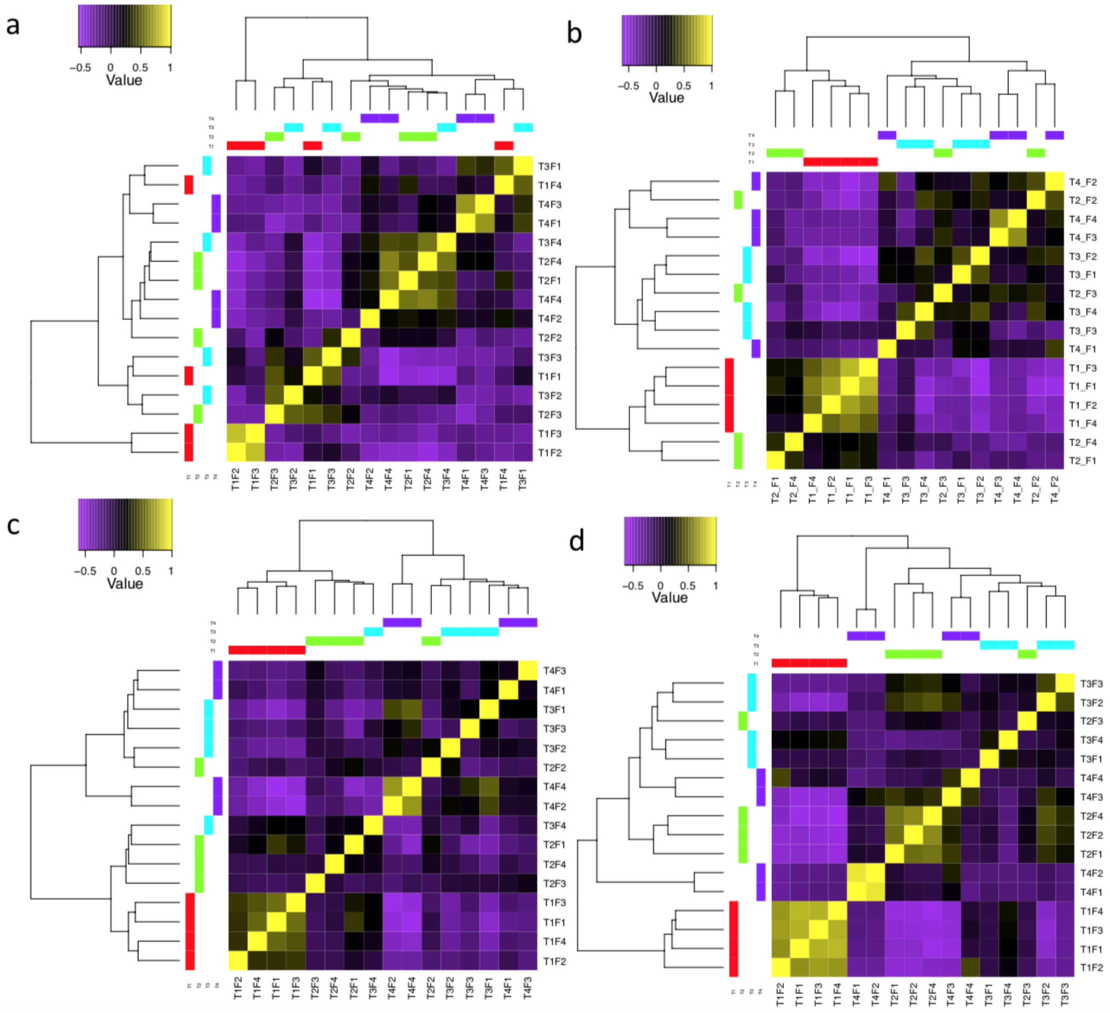


**Supplementary Figure S2. Variation among the four biological replicates at the four sampling events (T1, T2, T3 and T4).** The heatmaps show the hierarchical clustered Spearman correlation resulting from comparison of genome-wide expression values ($log_2$CPM) for all samples against each other in brain **a**, pituitary **b**, ovary **c**, and liver **d**. The level of correlation is presented by a colour field. Sample clustering revealed grouping of the samples from mature fish specially at T3 and T4 in pituitary, ovary and liver. Brain samples revealed high variability among biological replicates within timepoint, likely due to inconsistent sampling of the heterogenous brain tissue. Brain samples were excluded from further analyses.

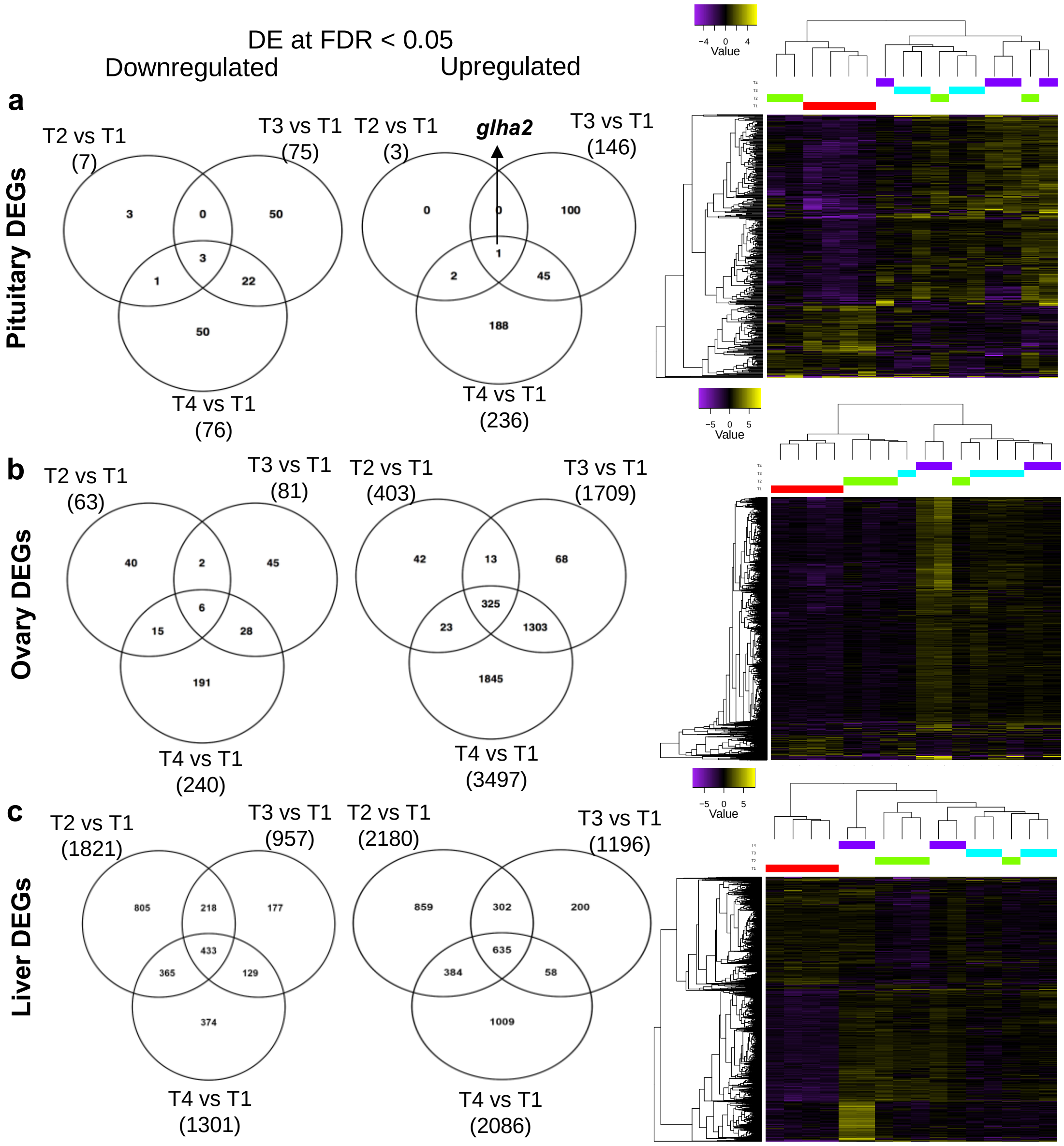


**Supplementary Figure S3. Gene expression in three tissues the during onset of Atlantic salmon maturation**. An adjusted $P < 0.05$ and logFC > ±1 was used to identify significantly differentially expressed genes (DEGs). The overlap between down- and up-regulated genes along with hierarchical clustering are shown for **a** pituitary, **b** ovary and **c** liver. It's worth noting that *glha2* (the common subunit present in gonadotropins) was consistently upregulated in the pituitary throughout the experiment. The clustering was obtained by comparing normalised expression (FPKM) for samples at T2, T3 and T4 in comparison to the control at T1. Expression values were $\log_2$-transformed and mean centred by gene. The relative expression values are shown in yellow-navy scale.

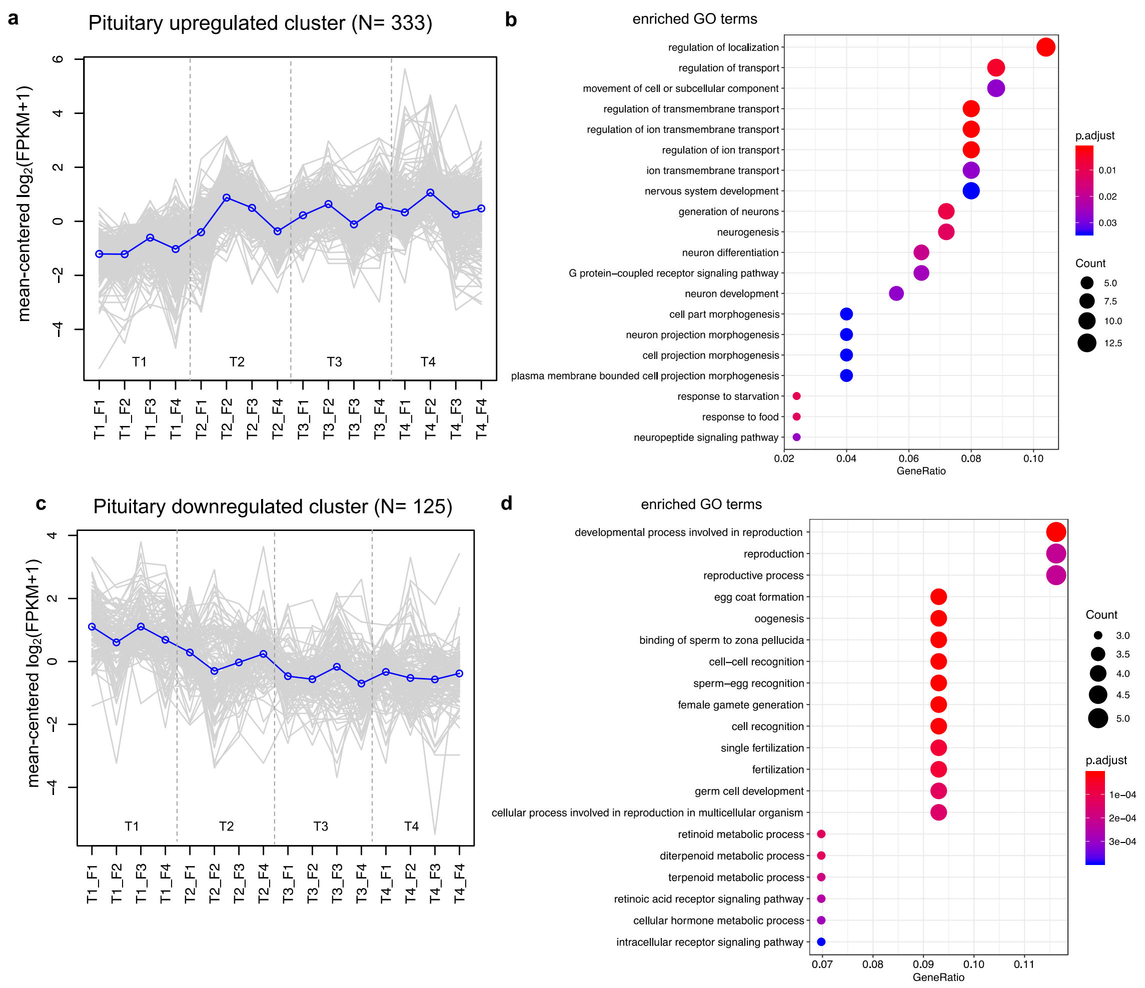


**Supplementary Figure S4. Gene cluster expression and GO enrichment for the pituitary.** The behaviour of genes identified in the upregulated and downregulated gene clusters described in **Supplementary Figure S3** are shown separately in **a** and **c**. The Y-axis represents the mean-centred $\log_2$(FPKM+1) value, expression of single genes is plotted in grey while the mean expression of the genes is plotted in blue. **b**, **d** Enriched gene ontology (GO) terms (hypergeometric test, Bonferroni-adjusted $P < 0.05$) among the list of the up- and (N=333) downregulated (N=125) and the gene ratio for the genes that map to each term.

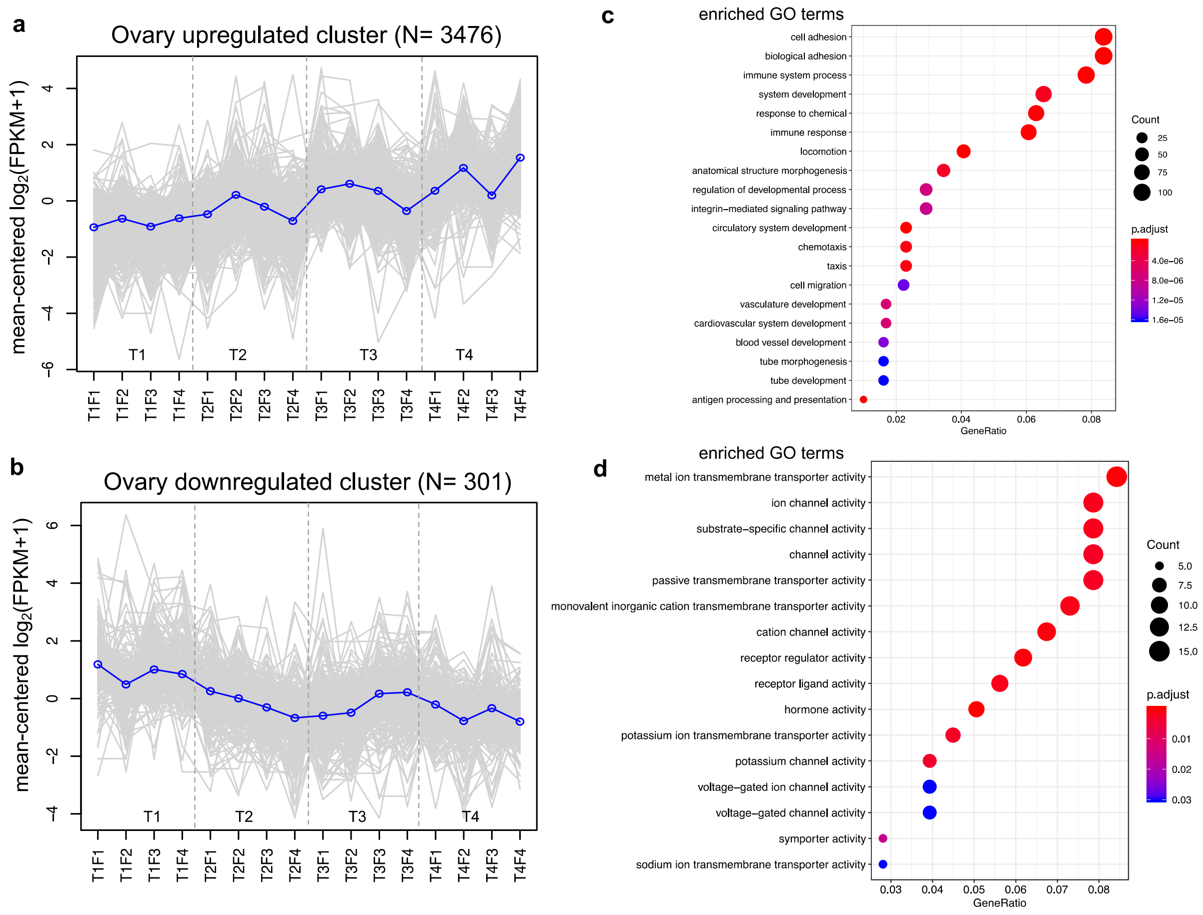


**Supplementary Figure S5. Gene cluster expression and GO enrichment for ovary**. The behaviour of genes identified in the upregulated and downregulated gene clusters described in **Supplementary Figure S3** are shown separately in **a** and **c.** The Y-axis represents the mean-centred $\log_2$(FPKM+1) value, expression of single genes is plotted in grey while the mean expression of the genes is plotted in blue. **c**, **d** Enriched gene ontology (GO) terms for each cluster (hypergeometric test, Bonferroni-adjusted $P < 0.05$ and the gene ratio for the genes that map to each term.

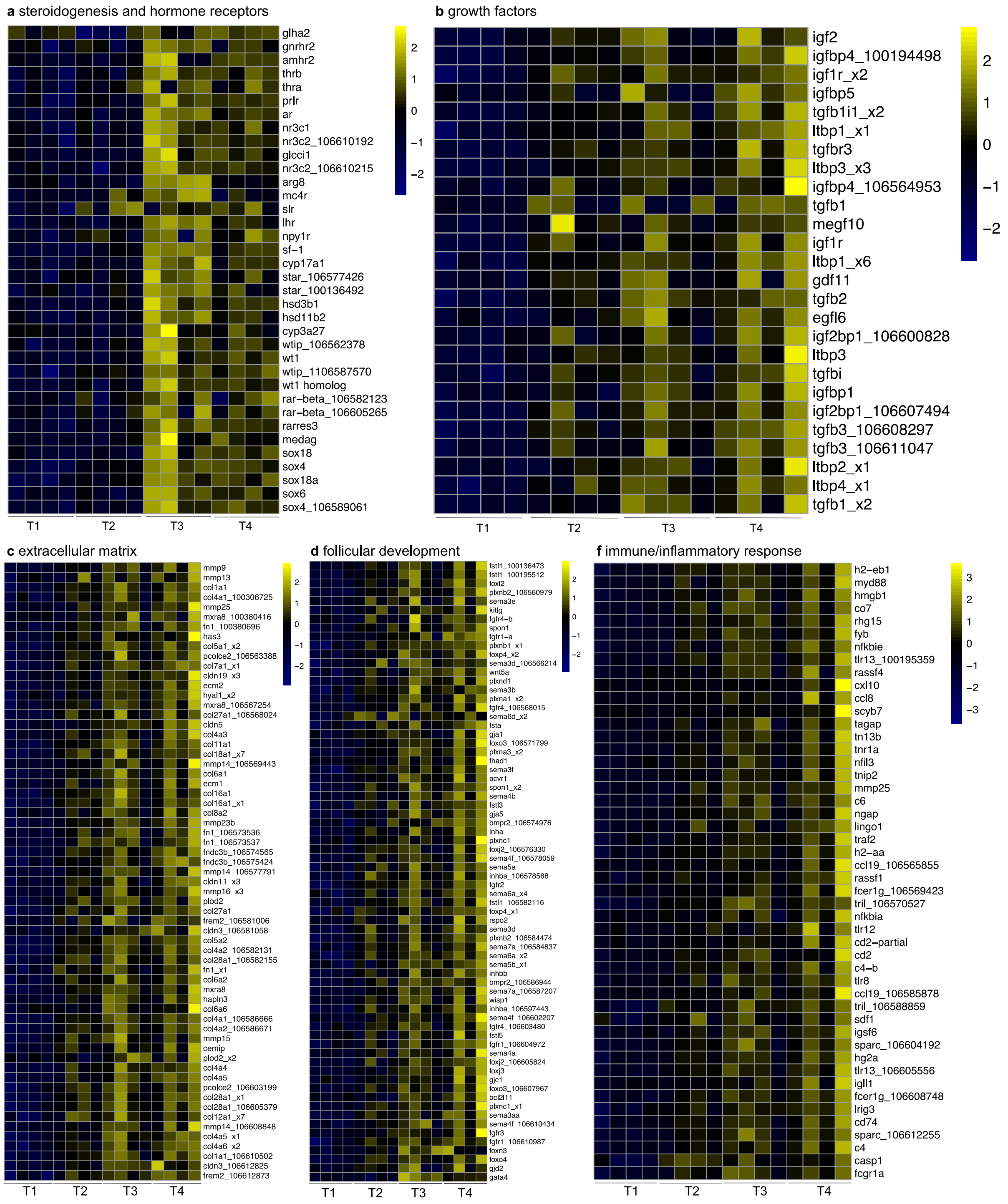


**Supplementary Figure S6. Ovary DEGs implicated in onset of salmon maturation**. The multi-panel heat maps show genes implicated in and/ or encoding **a**, steroidogenesis and hormone receptors. **b**, growth factors. **c**, extracellular matrix remodelling. **d**, follicular development. **e**, immune- and inflammatory responses. The clustering shown was obtained by comparing normalised expression (FPKM) for samples at T2, T3 and T4 compared to the control at T1. Expression values were $log_2$-transformed and mean centred by gene. The relative expression values are shown in yellow-navy scale.

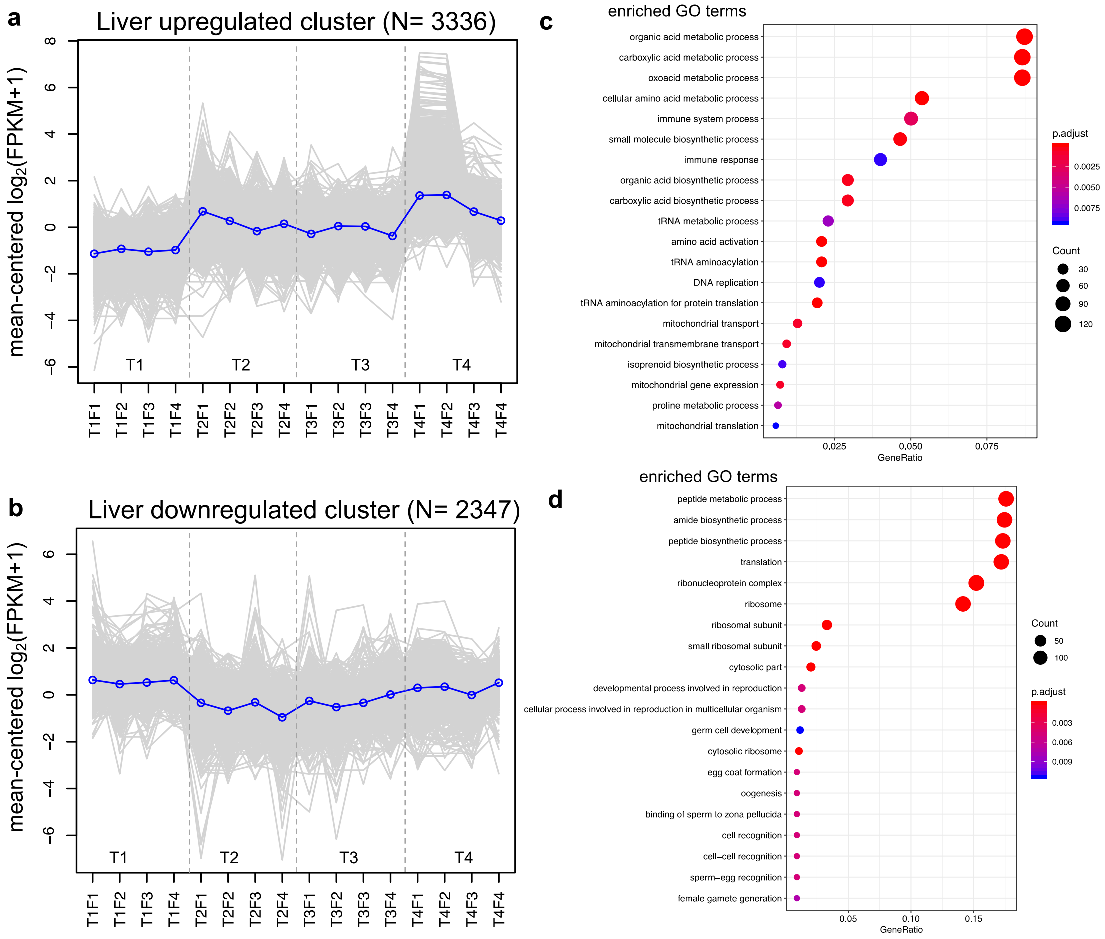


**Supplementary Figure S7. Gene expression and GO enrichment for the liver.** The behaviour of genes identified in the upregulated and downregulated gene clusters described in **Supplementary Figure S3** are shown separately in **a** and **c**. The Y-axis represents the mean-centred $\log_2$(FPKM+1) value, expression of single genes is plotted in grey while the mean expression of the genes is plotted in blue. **c**, **d** Enriched gene ontology (GO) terms for each cluster (hypergeometric test, Bonferroni-adjusted P < 0.05 and the gene ratio for the genes that map to each term.

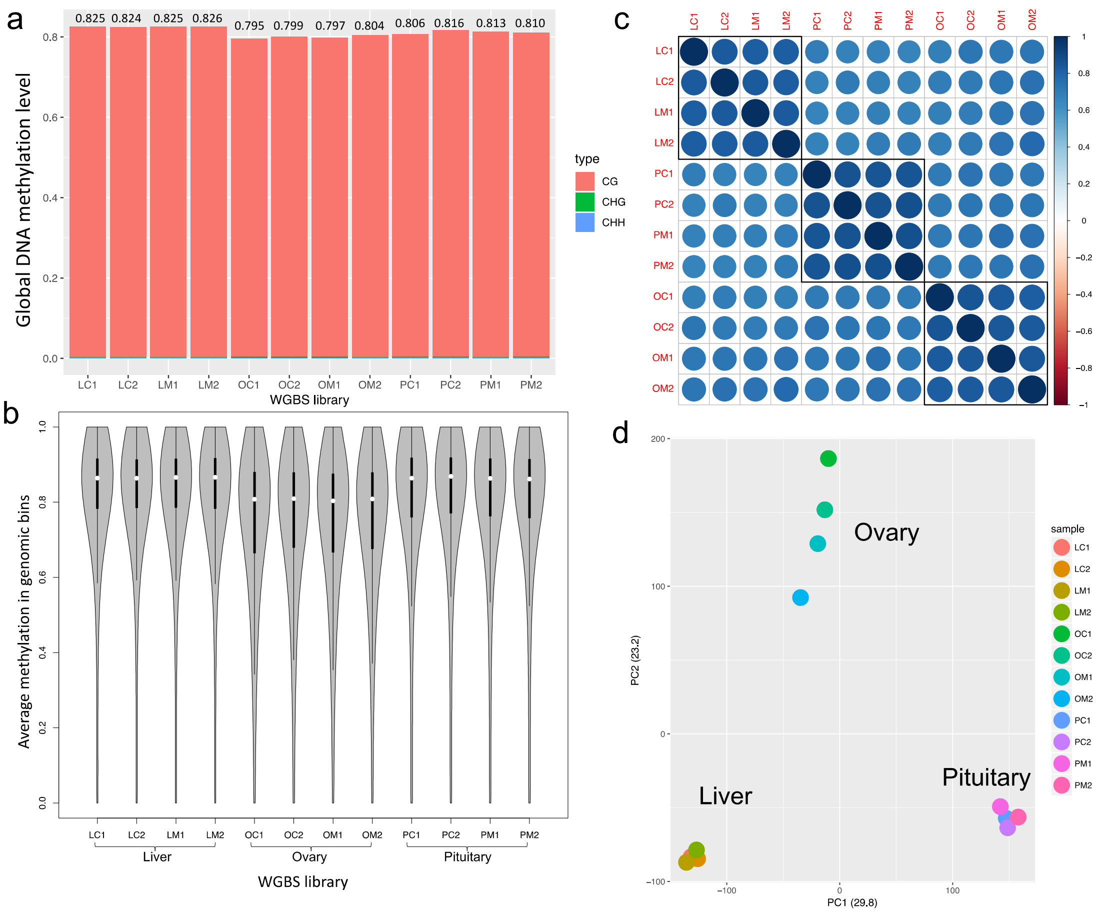


**Supplementary Figure S8. Characterization DNA methylomes of three Atlantic salmon tissues**. **a**, Global DNA methylation levels at different CG context (CG, CHG and CHH) in liver (L), ovary (O) and pituitary (P). Samples are named as either C (for control at timepoint 1) or M (for mature at timepoint 4) and with a number (1 or 2) to distinguish biological replicates. CpG methylation contributes to 99.5% of methylated Cs. **b**, Violin plot showing average CpG methylation levels in genomics bins across the twelve methylome libraries shows similar profiles for samples of each tissue. **c**, Correlation matrix of replicates using common CpGs, replicates of the same tissue shows higher correlation and clustered together. **d**, Principal Component Analysis (PCA) using common CpGs separates samples by tissues along PC1 (explaining 29.8% of the variance) and shows extensive variation among ovary methylomes along PC2.

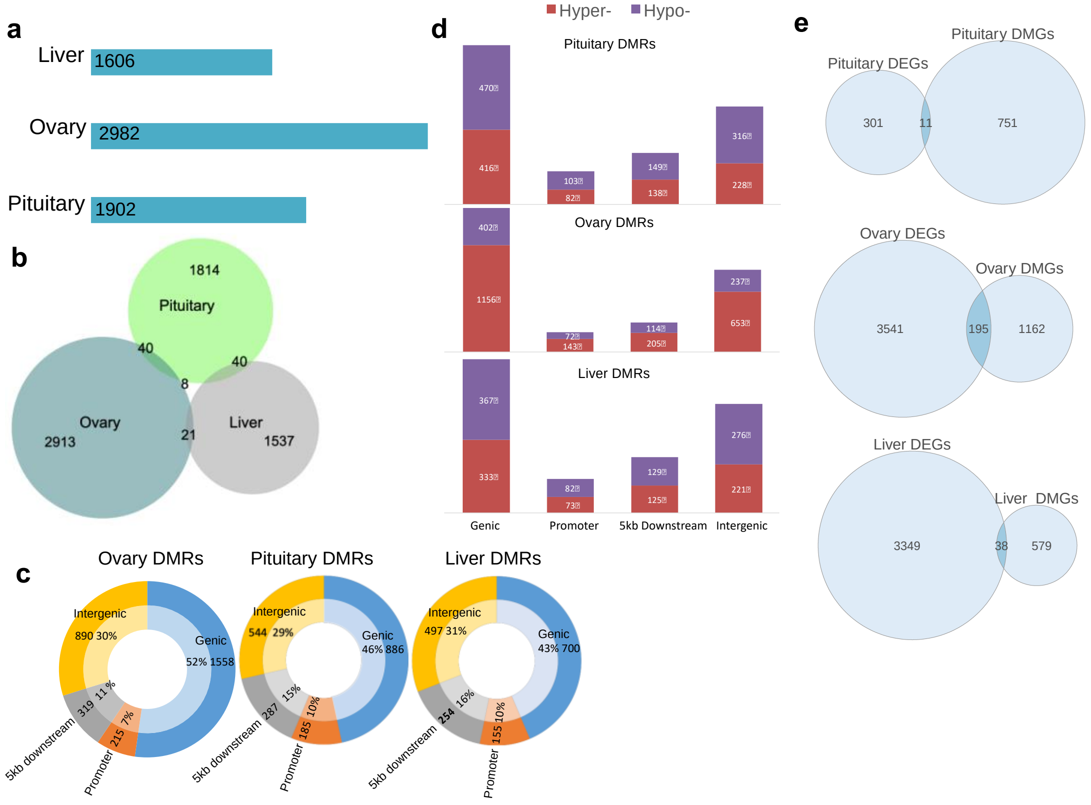


**Supplementary Figure S9. Multi-tissue genome-wide differences in CpG methylation during onset of salmon maturation**. A *P* threshold of 0.001 and delta (difference of the two group means) > 10% were used to identify significantly differentially methylated regions of the genome (DMRs). **a**, Bar graph showing the number of declared DMRs for each tissue. **b**, Venn diagram shows the overlap of DMRs across the three tissues. **c,** Relative distribution of DMRs across genic, promoters, 5kb downstream and intergenic regions of the Atlantic salmon genome in pituitary, ovary and liver are shown. DMR genomic distribution reveals on average 70% of DMRs were co-located within 5kb of a gene. **d**, The direction of methylation changes (hyper- and hypomethylation) across the four genomic categories in the three tissues. **e**, Overlap between sets of DMGs and DEGs detected at T4 vs T1 in the three tissues.

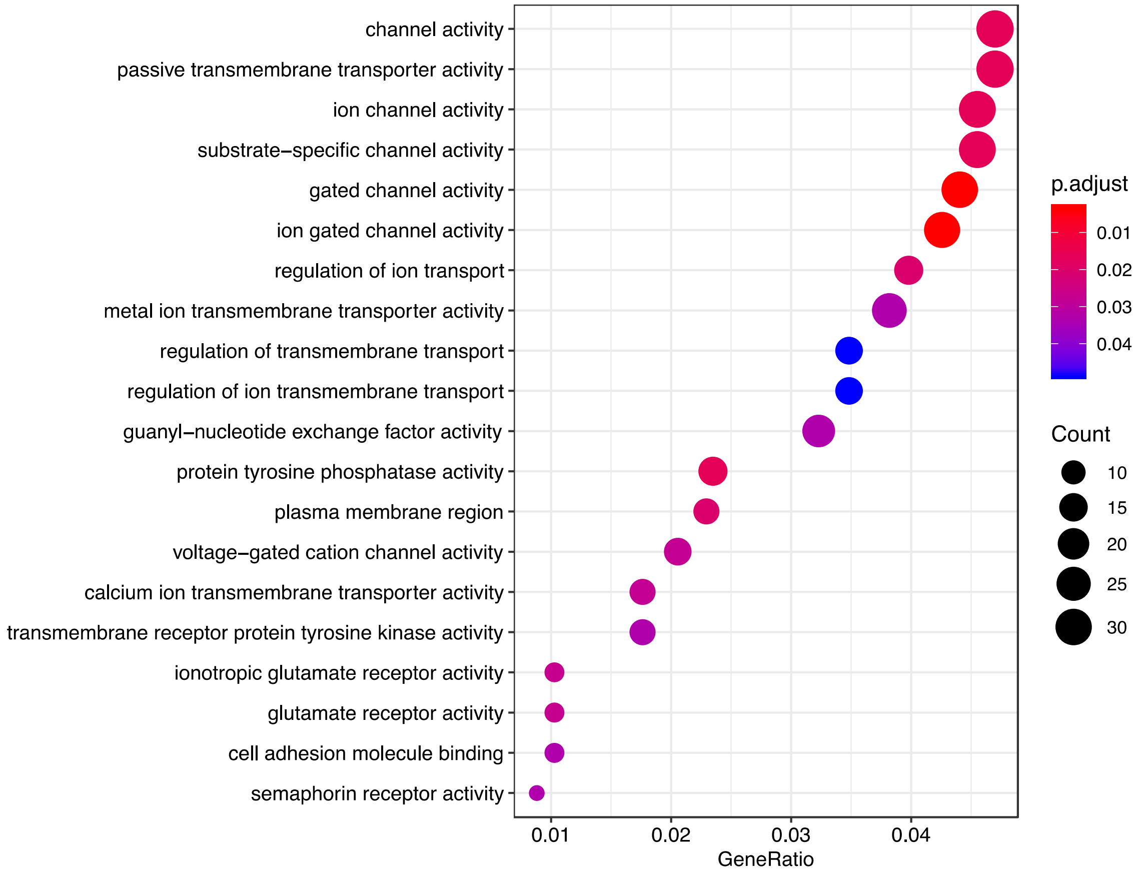


**Supplementary Figure S10**. **Enriched gene ontology (GO) terms among ovary hypermethylated genes** (n= 1156) against background of ovary genes (hypergeometric test, Bonferroni-adjusted *P* < 0.05) and the gene ratio for the genes that map to each term.

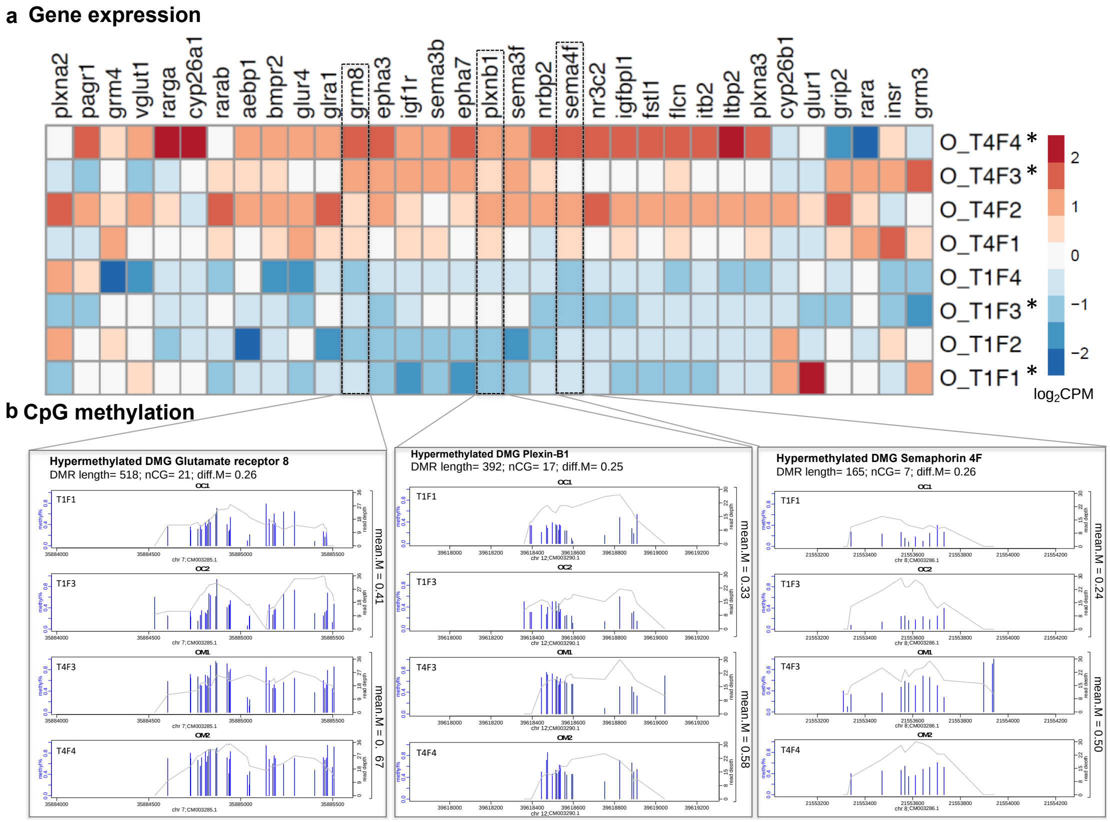


**Supplementary Figure S11. DNA methylation and expression patterns of biological candidates. a**, The expression profile of 33 biologically relevant hypermethylated genes in ovary. The $log_2$FPKM data was plotted separately for each of the four replicates at both T1 and T4. Stars highlight the four samples used for generation of the methylome data. **b** Patterns of CpG methylation for three candidate hypermethylated genes in ovary. The plot shows methylation percentages along with coverage depth at each CpG site. Pink rectangles represent the differentially methylated regions between two replicates of T1 (top 2 panels) and T4 (bottom 2 panels). Genomic coordinates are indicated below each density plots.

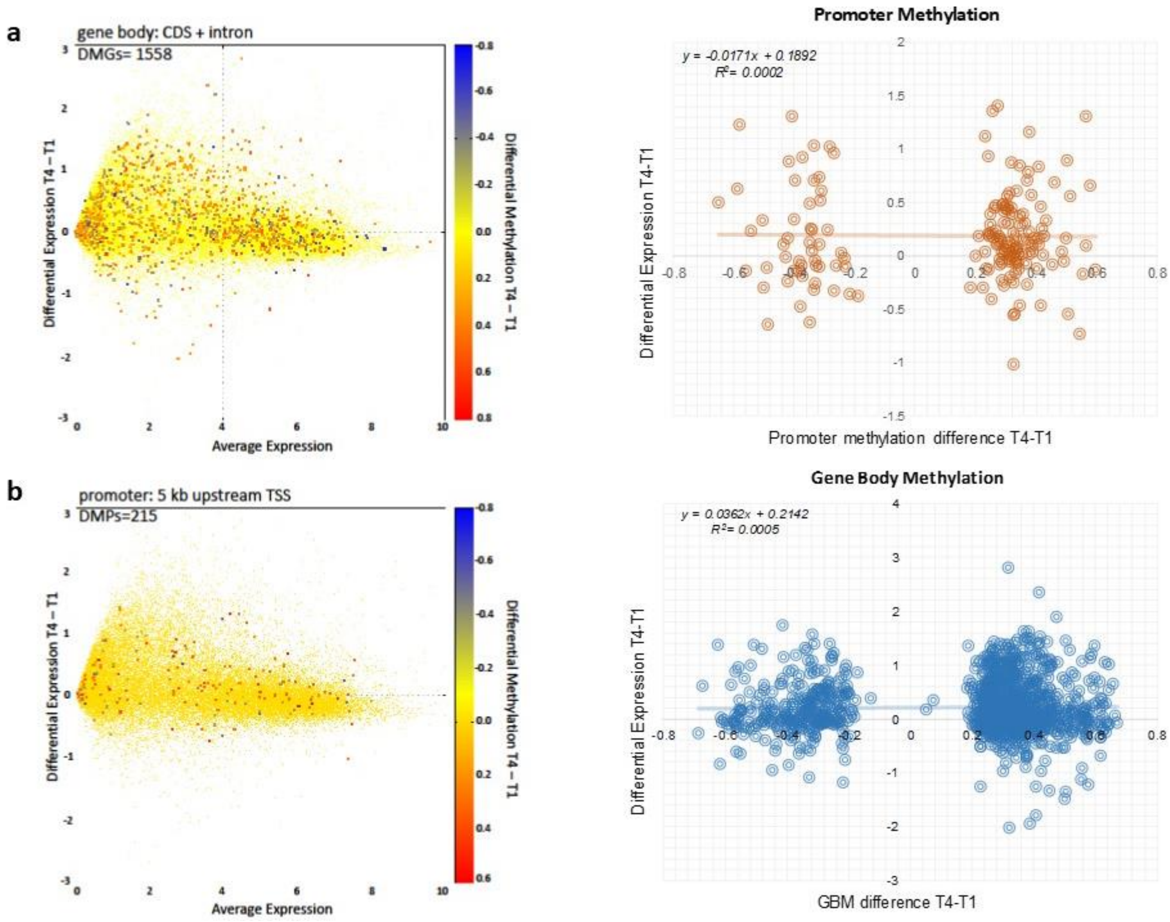


**Supplementary Figure S12. Correlation between changes in DNA methylation and gene expression in ovary**. The MA biplots show an absence of correlation between genome-wide differential gene expression in T4-T1 ovary samples in yellow and methylation levels in blue-red spectrum reflecting hypo-to hypermethylation at both gene bodies (**a**) and promotor regions (**b**). Linear regression analyses further confirmed very low R-squared values in both cases.

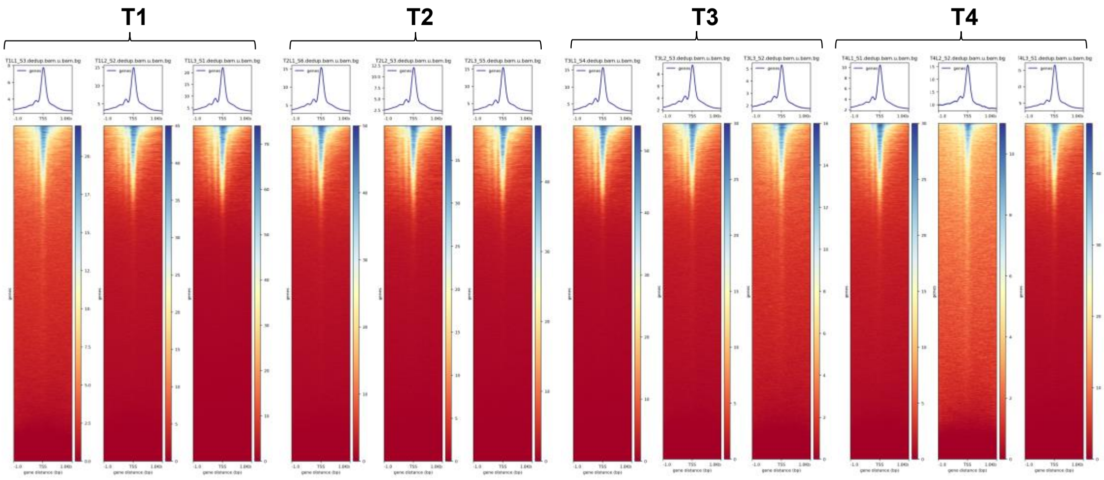


**Supplementary Figure S13. ATAC-seq library quality control.** Evaluation of ATAC-seq libraries using transcription start site enrichment (TSSs) analysis. The plots show density plots and heatmaps for reads ±1 kb around TSSs for 12 ATAC-seq liver libraries (4 time points x 3 replicates).

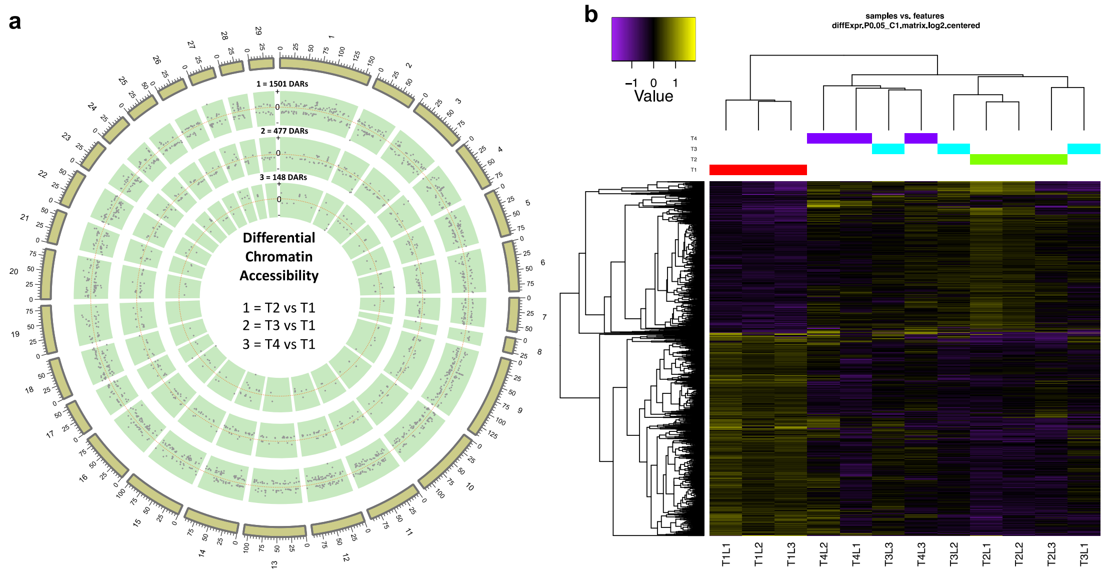


**Supplementary Figure S14. Changes in chromatin accessibility in liver during maturation onset. a** The circos plot shows the genomic distribution of the differentially accessible regions (DARs) (FDR < 0.05 and log2FC > ±1) detected in the triplicates of T2, T3, T4 samples (from outer circle inwards) compared to the control T1. **b** Hierarchical clustering of 1,831 unique DARs detected across time points in liver. The clustering was obtained by comparing normalised accessibility (CPM data) for samples at T2, T3 and T4 compared to the control at T1. Expression values were $log_2$-transformed and mean centred by peak. The relative accessibility values are shown in yellow-navy scale.

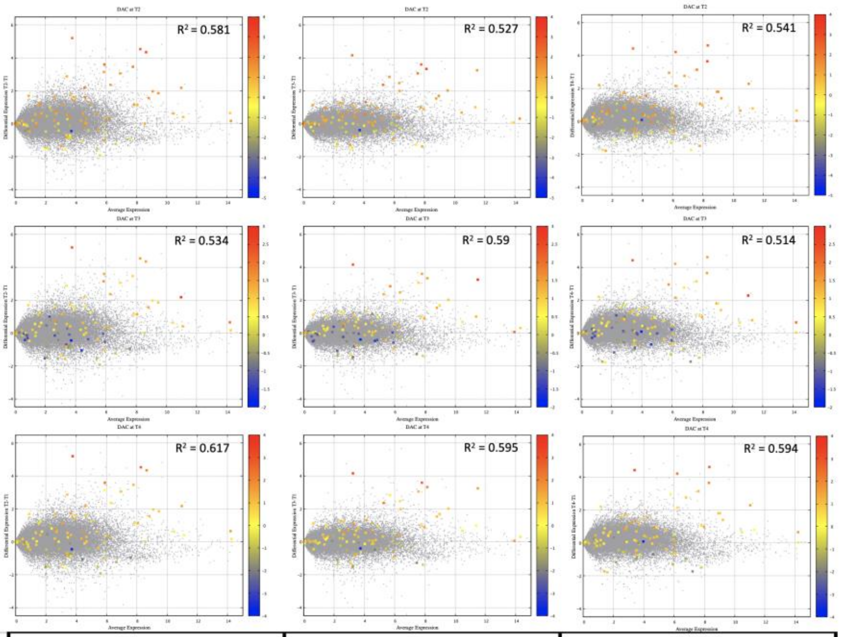


| Proximal DACs | DAC T2 | | | DAC T3 | | | DAC T4 | | |
|---|---|---|---|---|---|---|---|---|---|
| | b | b2 | R2 (%) | b | b2 | R2 (%) | b | b2 | R2 (%) |
| **DEG T2** | 0.63817 | 0.21864 | 46.3 | 0.65643 | 0.21117 | 31.6 | 0.71544 | 0.1949 | 46.4 |
| **DEG T3** | 0.47071 | 0.166 | 38.3 | 0.56949 | 0.23685 | 40.7 | 0.5603 | 0.13725 | 41.7 |
| **DEG T4** | 0.52188 | 0.20852 | 40 | 0.54754 | 0.25599 | 31.9 | 0.62755 | 0.19439 | 45.3 |

**Supplementary Figure S15. Positive correlation between gene expression and chromatin state accessibility at promoters.** The 9 MA biplots show genome-wide gene expression and overlaid with chromatin state accessibility data for DARs at promoters in liver at T2, T3, T4 compared to T1. The table shows results of a regression analysis conducted on significant DARs located 5 kb upstream of transcription start sites (TSSs) and gene expression data of the nearest genes.

| Rank | Motif | P-value | log P-pvalue | % of Targets | % of Background | STD(Bg STD) | Best Match/Details |
|---|---|---|---|---|---|---|---|
| 1 | GTACAGAA | 1e-37 | -8.638e+01 | 57.63% | 3.19% | 56.3bp (30.8bp) | CNOT4(RRM)/Homo_sapiens-RNCMPT00156-PBM/HughesRNA(0.853) More Information \| Similar Motifs Found |
| 2 | TTGATGTG | 1e-30 | -7.126e+01 | 50.85% | 2.71% | 62.0bp (51.5bp) | PH0134.1_Pbx1/Jaspar(0.779) More Information \| Similar Motifs Found |
| 3 | TCAGCATG | 1e-29 | -6.764e+01 | 49.15% | 4.07% | 55.4bp (0.0bp) | Zic(Zf)/Cerebellum-ZIC1.2-ChIP-Seq(GSE60731)/Homer(0.801) More Information \| Similar Motifs Found |
| 4 | CCTATTTT | 1e-26 | -6.062e+01 | 45.76% | 5.22% | 54.3bp (64.7bp) | CPEB2(RRM)/Homo_sapiens-RNCMPT00012-PBM/HughesRNA(0.868) More Information \| Similar Motifs Found |
| 5 | ATCAGCCA | 1e-24 | -5.708e+01 | 72.88% | 14.65% | 56.4bp (47.9bp) | CRZ1(MacIsaac)/Yeast(0.780) More Information \| Similar Motifs Found |
| 6 | GAGTGACC | 1e-19 | -4.424e+01 | 37.29% | 3.57% | 49.8bp (0.0bp) | Unknown1(NR/Ini-like)/Drosophila-Promoters/Homer(0.818) More Information \| Similar Motifs Found |
| 7 | TGTTGACA | 1e-17 | -4.005e+01 | 44.07% | 6.35% | 56.4bp (45.2bp) | Tbx20(T-box)/Heart-Tbx20-ChIP-Seq(GSE29636)/Homer(0.868) More Information \| Similar Motifs Found |
| 8 | CATTGCAA | 1e-16 | -3.817e+01 | 33.90% | 2.18% | 60.2bp (0.0bp) | slbo/dmmpmm(Pollard)/fly(0.781) More Information \| Similar Motifs Found |
| 9 | GAAATCTT | 1e-14 | -3.266e+01 | 45.76% | 10.34% | 56.8bp (36.9bp) | PH0037.1_Hdx/Jaspar(0.744) More Information \| Similar Motifs Found |
| 10 | TGTCAAGT | 1e-14 | -3.241e+01 | 30.51% | 5.21% | 51.1bp (7.0bp) | vnd/dmmpmm(Noyes_hd)/fly(0.840) More Information \| Similar Motifs Found |
| 11 | GTGCCCAT | 1e-13 | -3.040e+01 | 44.07% | 10.15% | 51.2bp (3.0bp) | PB0133.1_Hic1_2/Jaspar(0.833) More Information \| Similar Motifs Found |
| 12 | ACTCAGTC | 1e-13 | -3.016e+01 | 54.24% | 13.34% | 57.5bp (39.5bp) | z/dmmpmm(Bigfoot)/fly(0.754) More Information \| Similar Motifs Found |
| 13 | ACACCCCC | 1e-12 | -2.964e+01 | 28.81% | 4.88% | 46.6bp (23.7bp) | Run/dmmpmm(Papatsenko)/fly(0.815) More Information \| Similar Motifs Found |

**Supplementary Figure S16. Motif Enrichment Analysis.** Known motif enrichment results obtained using Homer software ($P < 1e^{-10}$; at least 25% of targets) by comparing accessible regions at promoters in the accessible cluster to a background of inaccessible regions at promoters in the inaccessible cluster.

**a**

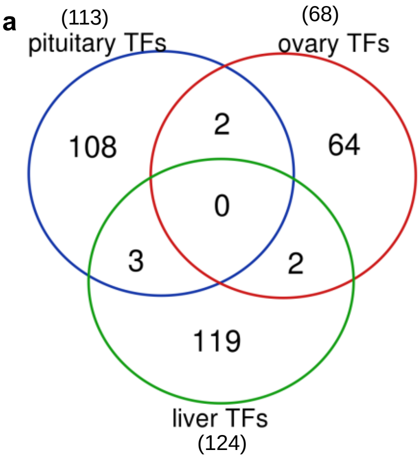


**b**

| overlap | number | geneID | description |
|---|---|---|---|
| pituitary-ovary | 2 | 106567572<br>106587538 | tubulin-specific chaperone cofactor E-like protein<br>LIM domain only 1 |
| ovary-liver | 2 | 106594966<br>106566725 | zinc finger protein 239-like<br>uncharacterized LOC106566725 |
| liver-pituitary | 3 | 106599217<br>106578771<br>106612998 | BTB/POZ domain-containing protein 6-B-like<br>AT-rich interactive domain-containing protein 4B-like<br>uncharacterized LOC106612998 |

**Supplementary Figure S17. Tissue overlap of transcription factors.** Overlap among transcription factors identified as master regulators according to the regulatory impact factor (RIF) metrics in pituitary and ovary and liver tissues ($P < 0.01$).

a

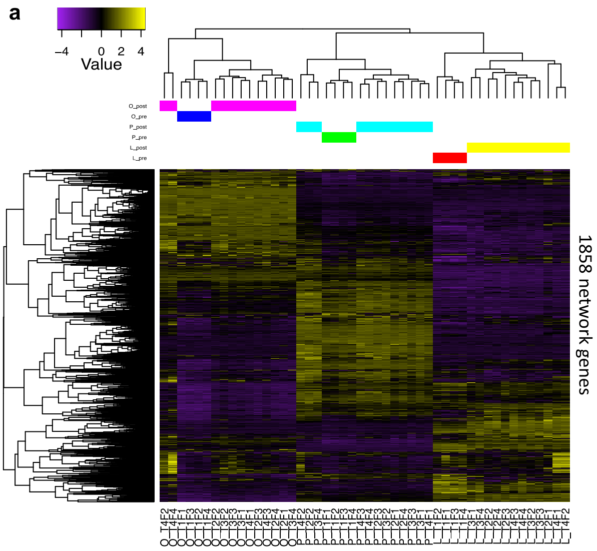


b

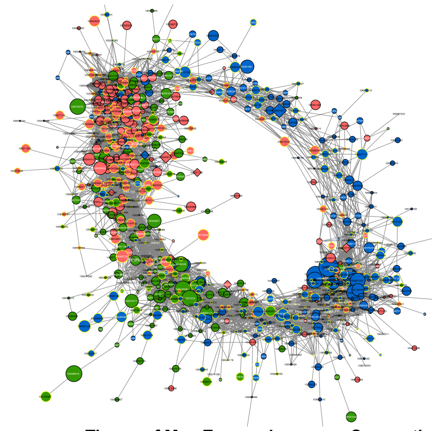

c

| | Tissue of Max Expression | | Connections | |
|---|---|---|---|---|
| **Tissue** | **N** | **%** | **N** | **%** |
| **Pituitary** | 777 | 41.82 | 353,004 | 42.27 |
| **Ovary** | 611 | 32.88 | 271,807 | 32.54 |
| **Liver** | 470 | 25.3 | 210,273 | 25.17 |
| **Total** | 1858 | 100 | 835,084 | 100 |

**Supplementary Figure S18. Multi-omics derived gene regulatory network. a** Heatmap of all 1,858 network genes. Genes (rows) and samples (columns) are organized by hierarchical clustering based on Euclidean distances. **b** Gene regulatory network constructed using the PCIT algorithm considering only genes with significant correlations ≥ ±0.95 (929 gene with 17708 connections). All nodes are represented by ellipses except for genes coding key regulators (TFs) have diamond shape. Nodes with yellow borders are differentially methylated, whereas nodes with white labels are differentially accessible. Node colours are relative to the tissue of maximum expression with blue represents the pituitary, red represents ovary and green represents liver. The size of the nodes is relative to the normalized mean expression values in all samples. **c** Tissue distribution and connectivity structures of the 1,858 network genes used to build the network.
